# Trait-dependent diversification in angiosperms: patterns, models and data

**DOI:** 10.1101/2022.05.18.490882

**Authors:** Andrew J. Helmstetter, Rosana Zenil-Ferguson, Hervé Sauquet, Sarah P. Otto, Marcos Méndez, Mario Vallejo-Marin, Jürg Schönenberger, Concetta Burgarella, Bruce Anderson, Hugo de Boer, Sylvain Glémin, Jos Käfer

## Abstract

Variation in species richness across the tree of life, accompanied by the incredible variety of ecological and morphological characteristics found in nature, has inspired many studies to link traits with species diversification. Angiosperms are a highly diverse group that has fundamentally shaped life on earth since the Cretaceous, and illustrate how species diversification affects ecosystem functioning. Numerous traits and processes have been linked to differences in species richness within this group, but we know little about how these interact and their relative importance. Here, we synthesized data from 152 studies that used state-dependent speciation and extinction (SSE) models on angiosperm clades. Intrinsic traits related to reproduction and morphology were often linked to diversification but a set of universal drivers did not emerge as traits did not have consistent effects across clades. Importantly, dataset properties were correlated to SSE model results - trees that were larger, older, or less well-sampled tended to yield trait-dependent outcomes. We compared these properties to recommendations for SSE model use and provide a set of best practices to follow when designing studies and reporting results. Finally, we argue that SSE model inferences should be considered in a larger context incorporating species’ ecology, demography and genetics.

## Introduction

Species diversity is unevenly distributed across the tree of life and while substantial research has investigated why some clades are more diverse than others, many fundamental questions remain unanswered. The causes behind this unevenness can be diverse, from catastrophic mass extinctions that decimate diversity (Raup & Sepkoski, 1982) to key innovations that spur on rapid speciation (Hodges & Arnold, 1995) to ecological factors such as competition that shapes species co-existence (Drury et al., 2016; Rabosky, 2013). A greater understanding of the drivers of species diversification is important because they can provide insights into the assembly of communities and their phylogenetic structure, the evolution of functional traits that underpin a species’ role in its environment, the formation of species interaction networks, and simply how biodiversity has evolved through time (Morlon, 2014). Research aiming to link species characteristics to macroevolutionary dynamics has been boosted over the last decade by the increasing availability of large phylogenetic trees (Jetz et al., 2012; Rabosky et al., 2018; Smith & Brown, 2018; Upham et al., 2019) and the continuing development of a range of statistical models to infer patterns in species diversification and the drivers behind them (Barido-Sottani et al., 2020; Beaulieu & O’Meara, 2016; Maliet et al., 2019; Rabosky & Huang, 2016). The increasing amount of empirical knowledge provides an opportunity to synthesise what we know so far about a wide range of ecologically diverse and species-rich clades to uncover general dynamics about the traits that have driven their diversification.

Species diversification can be linked to traits via state-dependent speciation and extinction (SSE) models. This popular family of models are based on birth-death processes where the diversification rates (birth is speciation, and death is extinction) are dependent on character states, and where transition rates between states define how state changes occur. The simplest SSE model is the binary-state speciation and extinction (BiSSE) model (Maddison et al., 2007) that takes as input a phylogenetic tree and state values (0 or 1) for each species in the tree. This allows users to uncover whether lineages with one state diversify faster than those with the other. SSE models can also be used to test whether the transition rates between states in one direction (0 to 1) are faster than the other (1 to 0). The original model has been extended in various ways (Fig. 1) to address different types of macroevolutionary questions. For example, ClaSSE (Goldberg & Igić, 2012) and BiSSE-ness (Magnuson-Ford & Otto, 2012) are extensions of BiSSE that include cladogenetic events (speciation simultaneously associated with change in state), and GeoSSE (Goldberg et al., 2011) explicitly models how diversification differs among geographic regions. Other developments include models with more than two character states (MuSSE; FitzJohn, 2012), quantitative traits (QuaSSE; FitzJohn, 2010) and semi-parametric models (FiSSE; Rabosky and Goldberg, 2017). Perhaps the most important innovation after the initial wave of SSE models was the introduction of hidden states into SSE models (Beaulieu & O’Meara, 2016; Caetano et al., 2018; Herrera-Alsina et al., 2019) as a way to account for background heterogeneity that can lead to false positives (Rabosky & Goldberg, 2015). The incorporation of hidden states into SSE models allowed diversification rates to be influenced by the focal traits as well as an unobserved trait(s) and provided a new set of more complex null hypotheses (the character independent (CID) models). This allowed users to test how relevant the focal trait is to species diversification (Beaulieu & O’Meara, 2016; Caetano et al., 2018) in the context of other factors. Here we focus on synthesizing results from SSE models used to investigate trait-dependent diversification in flowering plants, or angiosperms. There have been more than 150 such studies, providing an opportunity for an updated perspective on the role different traits have played in angiosperm diversification (Vamosi et al., 2018).

**Figure 1:**
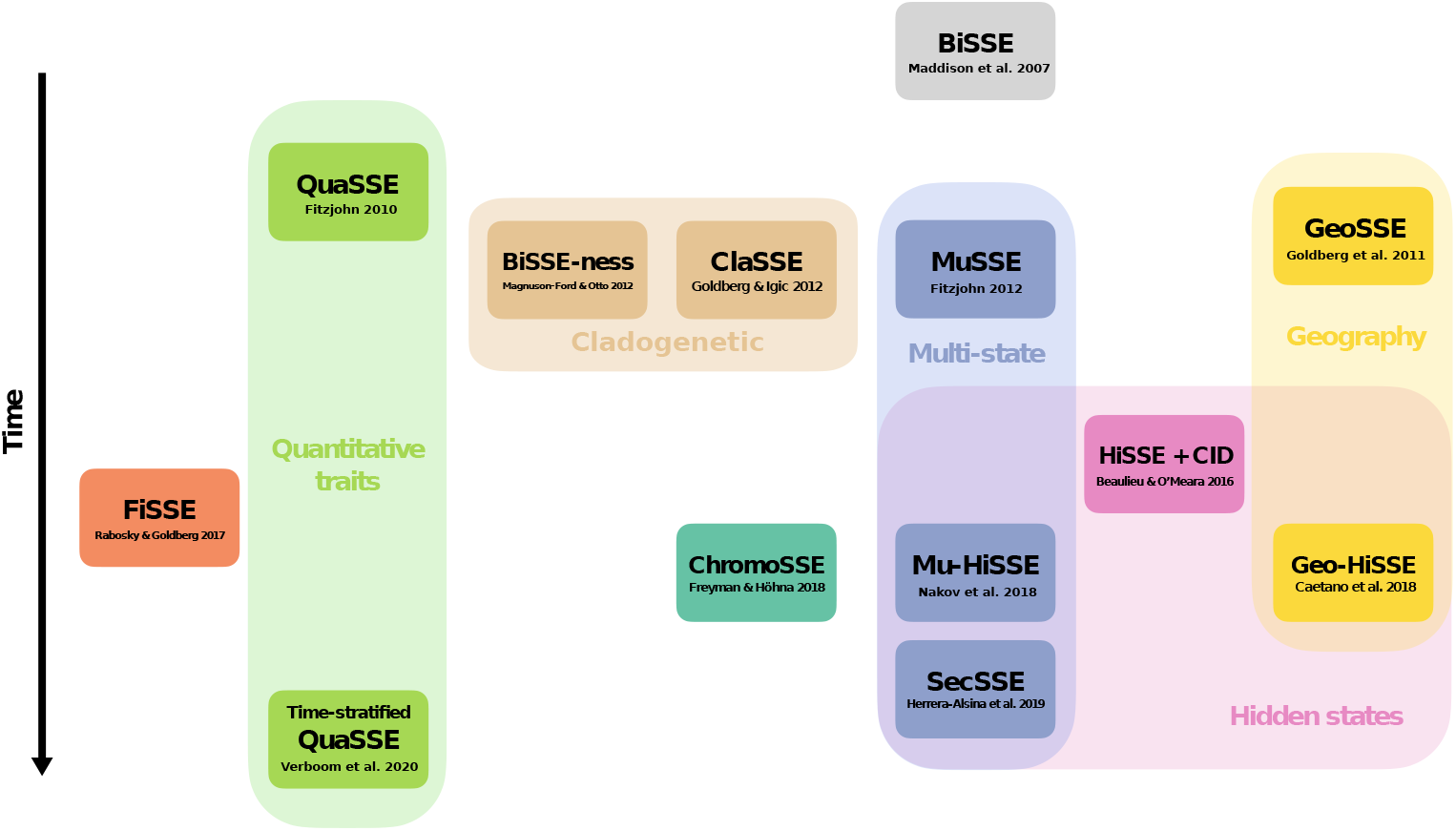
The development of state-dependent speciation and extinction (SSE) models. The original binary-state speciation and extinction model (BiSSE) model (Maddison et al., 2007) is shown at the top of the diagram with all other models depicted below, in the order of their publication. Acronyms are defined as follows; Binary-State Speciation and Extinction–node enhanced state shift (BiSSE-ness; Magnuson-Ford and Otto, 2012), Cladogenetic and Anagenetic Models of Chromosome Number Evolution (ChromoSSE; Freyman and Höhna, 2018),Character-Independent Diversification (CID Beaulieu and O’Meara, 2016), Cladogenetic State change Speciation and Extinction (ClaSSE; Goldberg and Igić, 2012), Fast, intuitive State-dependent Speciation-Extinction (FiSSE; Rabosky and Goldberg, 2017, Geographic State Speciation and Extinction (GeoSSE; Goldberg et al., 2011), Hidden Geographic State Speciation and Extinction (GeoHiSSE; Caetano et al., 2018), Hidden State Speciation and Extinction (HiSSE; Beaulieu and O’Meara, 2016), Multi-State Speciation and Extinction (MuSSE; FitzJohn, 2012), Multicharacter Hidden State Speciation and Extinction (Mu-HiSSE; Nakov et al., 2018), Quantitative State Speciation and Extinction (QuaSSE; FitzJohn, 2010; Verboom et al., 2020), Several examined and concealed states-dependent speciation and extinction (SecSSE Herrera-Alsina et al., 2019). Each box shows the name of the model and the associated citation. Models that share similar attributes (e.g. those with hidden states) are colour coded and grouped with boxes. This is not an exhaustive list of SSE models and does not include, for example, models used in epidemiology that allow tips to be sampled at various points in time (Scire et al., 2020).

Angiosperms form a clade of more than 350,000 extant species, despite their relatively young age of a 140 to 270 million years (Crepet & Niklas, 2009; Foster et al., 2017; Li et al., 2019; Magallón et al., 2015; Sauquet et al., 2021; Silvestro et al., 2021). Almost all of terrestrial life is linked, directly or indirectly to angiosperms (Benton et al., 2021) and their success makes them an ideal study group for uncovering the intrinsic traits and extrinsic factors driving their diversification. Previous work has suggested that the origins of angiosperm diversity can neither be tied to major global events nor the evolution of a single key innovation. Instead various combinations of traits, environment and ecology acting to stimulate diversification in different groups (Davies et al., 2004; Magallón & Castillo, 2009; Sauquet & Magallón, 2018), creating a landscape of macroevolutionary dynamics that vary substantially across different angiosperm clades (Magallón et al., 2019). One hypothesis proposes that the traits driving the differences in diversification are a range of vegetative and reproductive characteristics, some of which are unique to angiosperms (Stebbins, 1974). In sexual systems, for example, dioecy originated in 890-5000 independent instances (Renner, 2014) but these appear to have led to quite different macroevolutionary dynamics (Käfer et al., 2014; Sabath et al., 2016; Wang et al., 2021). In the majority of studied traits, we do not know how pervasive such differences are, nor have broad-scale empirical studies of trait-dependent diversification provided a general consensus on which traits are most important for angiosperm diversification.

In this study we bring together the latest empirical knowledge on angiosperm diversification to compare the effects of traits in different evolutionary contexts and identify those that have repeatedly stimulated diversification. We also investigate the relationship between the properties of datasets (e.g. tree size and global sampling fraction) and the results of published studies, highlighting how biases in our use of SSE models can affect our conclusions when searching for general trends. Finally, we identify gaps in our current knowledge and provide a set of best practices for diversification result-reporting to enhance our ability to fill these gaps in the future.

## Materials & Methods

### Data collection

We collected all published studies that cited SSE methods papers (Beaulieu & O’Meara, 2016; Caetano et al., 2018; FitzJohn, 2010, 2012; Freyman & Höhna, 2018; Goldberg & Igić, 2012; Goldberg et al., 2011; Herrera-Alsina et al., 2019; Maddison et al., 2007; Magnuson-Ford & Otto, 2012; Nakov et al., 2018; Rabosky & Goldberg, 2017; Verboom et al., 2020) using Google Scholar, last accessed 18th May 2021. To facilitate data collection from papers using SSE models, we developed a new R package called ‘papieRmache’ (https://github.com/ajhelmstetter/papieRmache). This package has two main purposes (1) to classify papers into different categories based on the frequency of term use in the text and (2) to pull out sections of the main text that contain a keyword or a pair of keywords while highlighting relevant information. We identified the SSE studies on angiosperms by using the keywords ‘angiosperm’, ‘flowering’ and ‘plant’ subsequently validating the subset of papers by hand and removing any studies on groups other than angiosperms. We then collected up to 30 different dataset properties from each paper relating to the trait investigated, the group studied, the phylogenetic tree and the outcome of the SSE model used (see appendix S1 for a detailed explanation of each property). In cases where there was uncertainty in how to interpret or collect data from a study we contacted the authors for their assistance and clarification, where possible.

### Trait classification

While some sets of character states were the same among studies (e.g. annual vs perennial; diploid vs polyploid), many of them did not overlap. We classified traits into different categories to facilitate comparisons among different trait types. At the broadest classification these were intrinsic (traits belonging to the species), extrinsic (environmental or geographic traits), interaction (traits related to other species), and combination (multiple traits belonging to different categories that were grouped, e.g. species that have both small fruits and are found on islands). To allow for analyses at different grouping levels we developed a trait ontology (Table S1) starting at the broadest level as just described and becoming more specific, up to level six.

### Data analysis

To examine the effect of particular traits on diversification we used the trait categories defined above and calculated the proportion of models in which trait-dependent diversification was inferred. In many cases multiple models are run per study, typically to investigate the effect of a single trait across different clades or the effects of different traits on diversification in a single clade. We considered each model separately here with an outcome of 1 (trait-dependent diversification detected) or 0 (no effect of trait detected) recovered per model. We examined patterns at different levels of trait categorization, as well as using only those models with hidden states. Whether or not trait-dependent diversification was detected was typically based on significance in model comparisons and/or posterior distributions of rates among states. However, if significance wasn’t inferred or reported, we followed the study narrative and statements made in the text. If model comparisons were conducted and reported, only the best-fitting model was considered, unless other models were explicitly referred to in the study. To facilitate comparison, we mainly consider whether or not a trait has an effect on diversification, irrespective of the direction of the effect (i.e. increase or decrease of diversification), as the direction is only defined at the state level. When both the effect and the absence of an effect of a trait on diversification has been inferred in different studies, we consider these studies ‘inconsistent’. Note that inconsistency doesn’t necessarily imply a contradiction, as the differences in the results of SSE models might be caused by differences in statistical power, type I errors, or biological differences between clades, as we will discuss below.

In models where information was available, net diversification rates (lineages per million years) were extracted for each character state. At a broad scale, relative differences in net diversification rates were calculated as (*r*_*max*_ − *r*_*min*_)*/r*_*max*_ and were used to represent the magnitude of the effect of a given trait on diversification, while taking into account general variation in diversification rates among clades. Comparisons were then made across trait level 1 and 2 categories. For ease of interpretation, these analyses were restricted to those models where all net diversification rates were positive. At a narrower scale, we identified eight traits for which there was enough replication to be able to assess whether one character state was consistently inferred to have higher net diversification rates than the other(s). We ensured these traits had been tested at least five times, in at least two different studies and two different clades.

We examined the relationship between SSE model inferences and continuous dataset properties; number of tips, root age, number of genetic markers, sampling fraction (here referring to global sampling fraction unless stated otherwise) and tip bias (here calculated as the number of tips with the most common state divided by the number of tips with rarest state). For each of these we constructed two density plots representing the distributions of values in cases where trait-dependent diversification was, and was not, inferred and compared the overlap between densities. We also fitted generalized additive models (GAM, Hastie and Tibshirani, 2017) to the continuous dataset properties with the SSE model result as a binary response variable (trait-dependent diversification vs no effect). The GAM approach allows linear or non-linear smooth functions to be used for predictor variables, giving greater flexibility in the estimation of relationships between predictors and the dependent variable. When analysing continuous data, all variables were log-transformed (or arcsine in the case of sampling fraction) to conform better to a normal distribution. Initially, we constructed a GAM using all five variables and assigned the mean of the known values to any missing values. We also assessed each variable individually to determine the shape of each relationship when examined in isolation. In all cases we used smoothing functions (cubic regression splines) and the dimension of the basis used to represent the smooth term was set to *k* = 5 for each variable.

### Predicting results based on dataset properties

After collecting information from all studies we found that the dataset properties were sometimes associated with the outcome of the SSE model, that is, whether trait-dependent diversification was inferred or not. We therefore attempted to predict SSE model results (inference of trait-dependent diversification vs no effect) from dataset properties alone, and identify those properties with the largest predictive power. We used all available dataset properties except for highly-specific categorical variables (e.g. trait levels 5-6, clade, family) and those that varied among different states (putative root state, sampling per state, samples per state). We used a machine learning approach, extreme gradient boosting, with the R package ‘xgboost’ (Chen & Guestrin, 2016), a supervised learning approach based on gradient boosting machines. This family of methods uses a labelled dataset (the outcome is known) and an ensemble of weak prediction models (e.g. decision trees) whereby new models are added on to existing models per iteration to minimize error. The xgboost algorithm improves upon other boosting methods with its increased speed and enhanced regularization to minimize overfitting (Chen & Guestrin, 2016). Prior to running our models, categorical variables with more than two categories were converted into binary, dummy variables using one-hot encoding (i.e. each unique category is converted into its own binary variable) to facilitate model building. We trained models on a random selection of 80% of our dataset and tested them on the other 20%. After a parameter optimisation step we repeated this process 500 times to produce a range of accuracy values, the percentage of cases where the real outcome matched the classification, to account for stochasticity in the test and training datasets. For each iteration we also recovered the relative importance of each variable, which allowed us to determine which dataset properties had the most influence on the model. Given the inter-dependency of variables across the different decision trees, it is difficult to uncover whether a given property generally leads to trait-dependent diversification or not with xgboost. We avoid interpreting the results in this way, focusing on how accurate prediction can be and the variables that are most important to the model’s predictive ability.

## Results

### Traits studied and their effects on diversification

We collated information on trait-based diversification from 152 studies using a total of 629 SSE models to study angiosperm diversification. We found that 124 studies were conducted on a single clade, the rest examined diversification patterns across multiple clades. Variation in breadth of different traits investigated was also observed within studies. In total, 92 studies examined just a single trait (i.e. one level 6 category, see Table S1), while 38 studies looked at diversification patterns in sets of traits that belonged to more than one trait category at the highest level (e.g. extrinsic and intrinsic traits). In terms of taxonomic level, SSE models were most often run on focal genera, or families (Fig. S1) and study clades were relatively evenly-distributed across the angiosperm tree of life (Fig. S2). Studies have focused on 36 out of 64 angiosperm orders, and 83 out of 416 families. As expected, diversification interest is generally proportional to the amount of species diversity in different parts of the angiosperm tree of life. There was a clear, positive correlation between the number of species in a clade (order, family) and the number of state-dependent diversification studies applied to the clade (Fig. S3, S4).

At the highest level of classification, intrinsic traits (i.e. those belonging to the plant species itself) were tested more often (295 models or 47% of models run) than extrinsic traits (i.e. those related to the species’ habitat and geography, 255 models or 41%). Researchers tended to study intrinsic traits relating to reproduction (e.g. flower morphology, fruit morphology, breeding system), traits related to species’ biogeography (e.g. biome, geographic region) and vegetative traits (e.g. life form, leaf morphology), investigating less often physiological characters (e.g. photosynthesis) or those related to interaction (e.g. symbiosis or dispersal) (Fig. 2). We compared the proportion of trait-dependent diversification outcomes in SSE models at different category levels. In general, intrinsic traits were found to be associated with diversification slightly more often than extrinsic traits (57.3% vs 52.5%).

**Figure 2:**
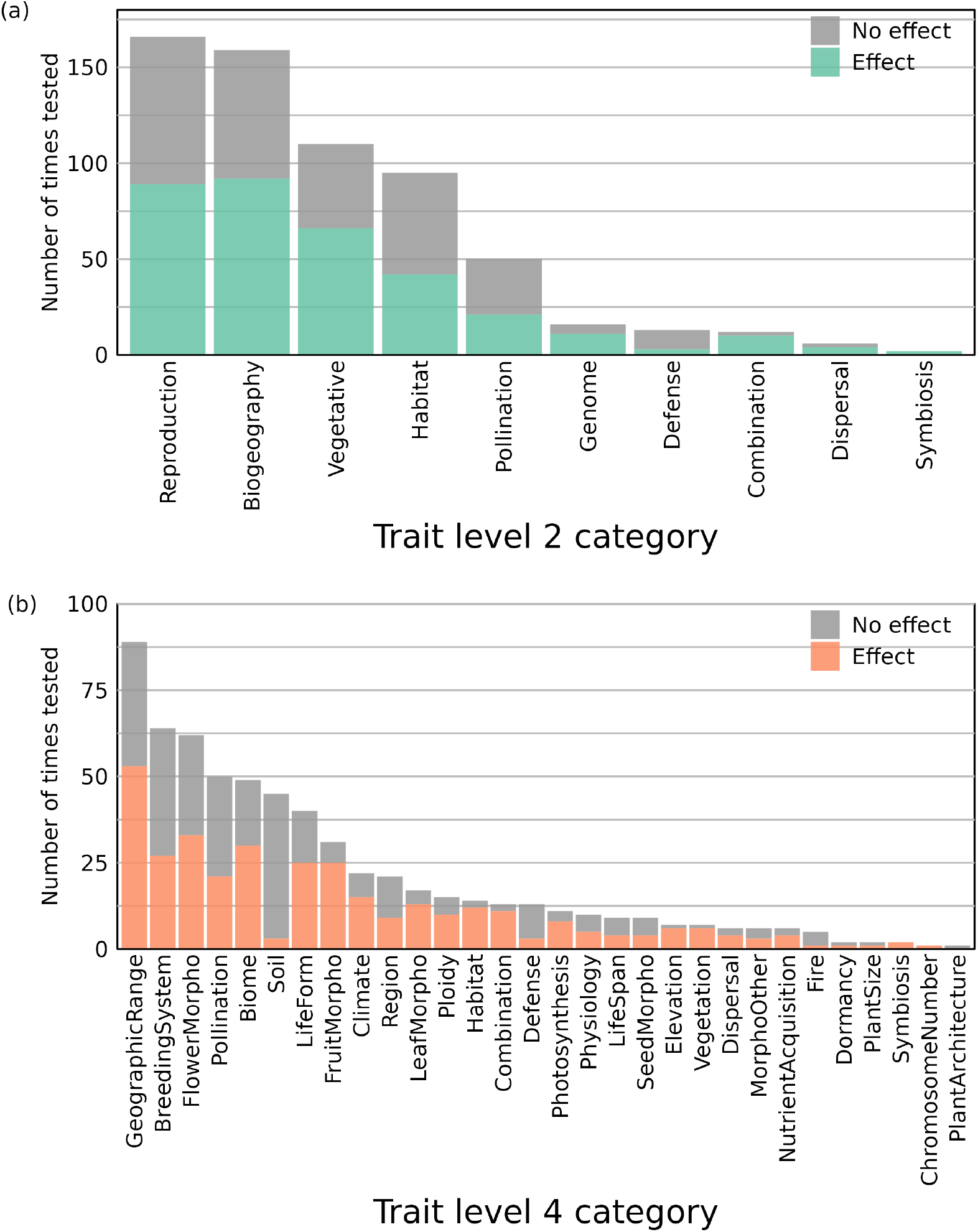
Stacked barplots showing how often particularly trait types were tested with state-dependent speciation and extinction (SSE) models. Bars are coloured to depict how often trait-dependent diversification was detected per trait type. If multiple SSE models were used in a single study they were considered separately i.e. each model contributed one result to the totals for each trait category. Two plots are shown, (a) one with relatively broad trait categories (level 2) and (b) one with narrower categories (level 4). An ontology depicting how different trait classification levels are connected can be found in Table S1.

If a trait has been studied more than once we can compare the effect of this trait on diversification in different evolutionary contexts to see if similar trends are found. Our collation of data showed that results inferred with SSE models were inconsistent at both broad and narrow scales (grey vs coloured portions of bars in Fig. 2). For example, traits such as lifespan (Azani et al., 2019; Drummond et al., 2012; Salariato et al., 2016; Soltis et al., 2013) and ploidy level (Folk & Freudenstein, 2014; Han et al., 2020; Landis et al., 2018; Zenil-Ferguson et al., 2019) yielded different results depending on the angiosperm group studied. Polyploidy has been linked to increased diversification in *Allium* (Han et al., 2020), while it had no effect on the diversification of Brassicaceae (Román-Palacios et al., 2019). Among those trait level 2 categories that have been tested using *>*25 models, vegetative traits yielded trait-dependent diversification more often than any other trait type, while pollination yielded the lowest proportion (Fig. 2a).

Though replication among character states was typically low we found eight traits that were tested often enough to assess whether there was a consistent effect of one state on diversification and the magnitude of the effect (Fig. S5). In three of these traits (lifespan, sexual system and woodiness) trait-dependent diversification was rarely found while in the remaining traits (epiphytism, biome, ploidy, photosynthesis and self-compatibility) results more often indicated trait-dependent diversification. However, we did find conflict in which states increased in diversification among different models in all traits except epiphyte form and self-incompatibility. Examining the absolute net diversification rates among states of seven traits (sexual system could not be assessed as most rates were not time-calibrated) we found that patterns across clades reflected those detailed above (Fig. S6). Net diversification rates in traits rarely associated with diversification (e.g. woodiness or lifespan) were generally similar among the different states (Fig. S6). To understand the effect of major trait categories on diversification we plotted the distribution of relative differences in net diversification rates for models belonging to each trait category (Fig. S7). Generally we found that there was a wide range of relative differences in each trait category but no statistically significant differences among categories.

### The evolution of SSE model use and methodological innovation

As SSE models themselves have diversified, the relative frequency of model-use has evolved. We collated data on the types of SSE model used in each study, and plotted their use by year of publication (Fig. 3). BiSSE has remained popular even as newer more complex models have emerged. Models with multiple states, predominantly MuSSE, have also been commonly used showing that researchers are interested in the effects of more complex traits or trait groups with more than two states. There has also been a consistent focus on using SSE approaches related to geography in models like GeoSSE and GeoHiSSE. When examining the number of studies that use SSE models each year we find a rapid increase since the first use of BiSSE on angiosperms in 2009 until a conspicuous slowdown and slight drop in 2015 (Fig. 3). This appears to coincide with the publication of a number of influential papers that criticised the propensity of SSE methods for false positives (Maddison & FitzJohn, 2015; Rabosky & Goldberg, 2015) and pointed out power limitations (Davis et al., 2013). After this, SSE model use continued with a greater variety of models owing to the development of models with hidden states (Beaulieu & O’Meara, 2016), which have since spread to all aspects of SSE model use (Fig. 1), becoming the dominant set of models by 2019 (Fig. 3). We tested whether the use of hidden state models (30 studies) lead to more consistent results than those reported for all studies (see above). We found that the proportion of trait-dependent outcomes increased for pollination, remained about the same for reproduction and decreased substantially for biogeography, vegetative and habitat (Fig. S8).

**Figure 3:**
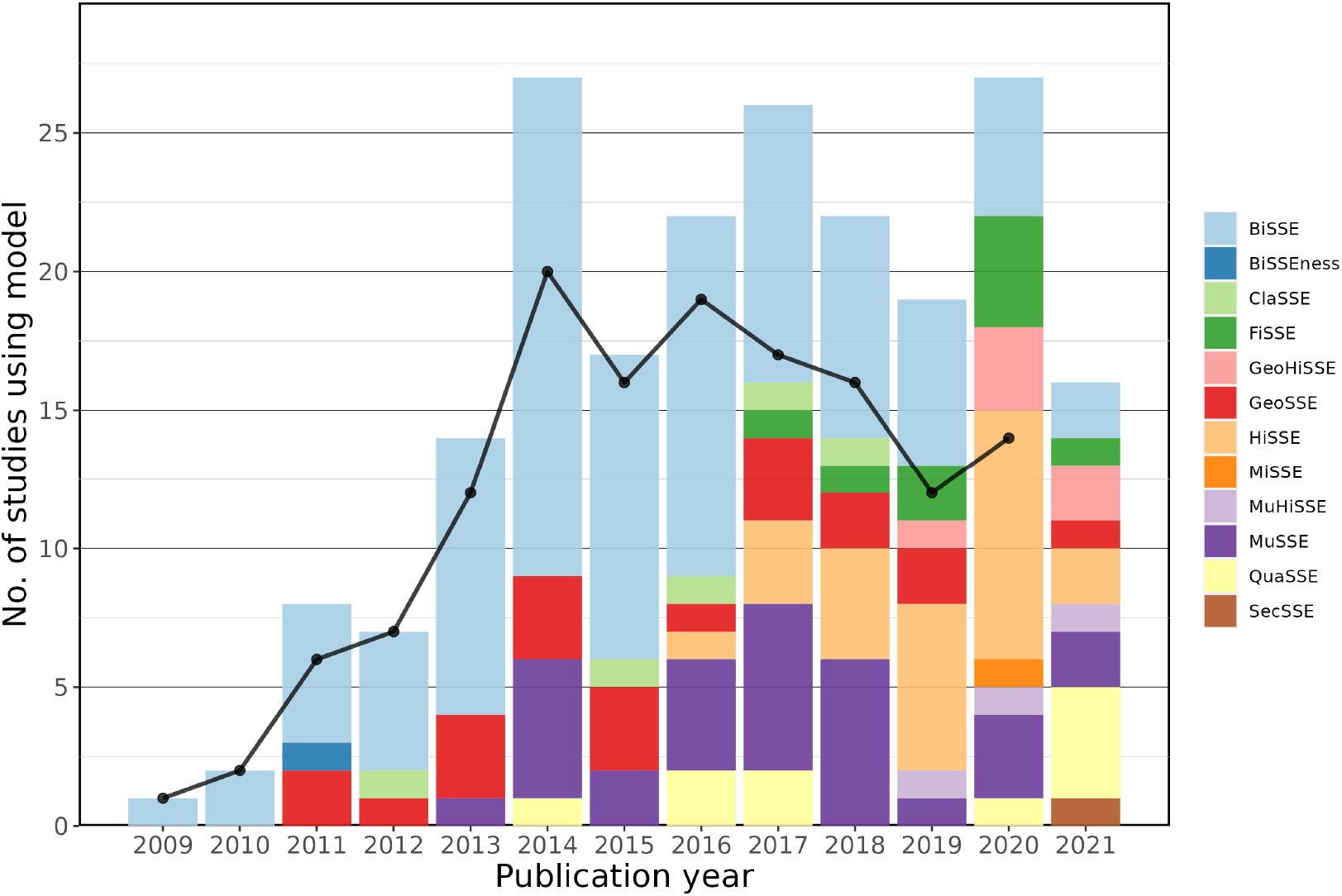
A stacked barplot showing the change in state-dependent speciation and extinction (SSE) models used on angiosperm clades over time. Each bar indicates the number of unique SSE model types per study totaled over the publication year. Bars are coloured according to the proportion of each SSE model type published in that year (see legend on the right of the plot). If the same SSE model was used multiple times in a single study it is only counted once (e.g. if BiSSE was used four times in a study published in 2012 this contributes an increase of one to the BiSSE portion of the 2012 bar). The black line shows the total number of studies using SSE models on angiosperms per year. Note that studies published after May 2021 were not included, so this year is incomplete.

### The importance of dataset properties

The input data for macroevolutionary studies have grown in size and quality, in parallel with the innovations in the SSE models. For example, we found evidence that over time, trees used with SSE models have gradually grown larger (Fig. S9). We examined the relationship between tree size and whether or not trait dependent diversification was inferred, regardless of the trait investigated. We found that, in general, trait-dependent diversification was detected less often when trees had smaller numbers of tips (Fig. 4a, S10a). The number of tips in a tree is important for robustness of SSE model results and guidelines for adequate power were put forward by Davis et al. (2013) who suggested that results from models using trees with fewer than 300 tips should be treated with caution. But has this recommendation shaped SSE model use? We examined sizes of trees used before and after this guideline was published, across all SSE models. The proportion of models run on trees with fewer than 300 tips was initially very high (94% of 139 total models) in studies published up until 2013. It then decreased to 57% (277 of 482) models in studies published from 2014 onwards. Despite this reduction, more than 60 models were run on trees with fewer than 50 tips after Davis et al. was published in 2013.

**Figure 4:**
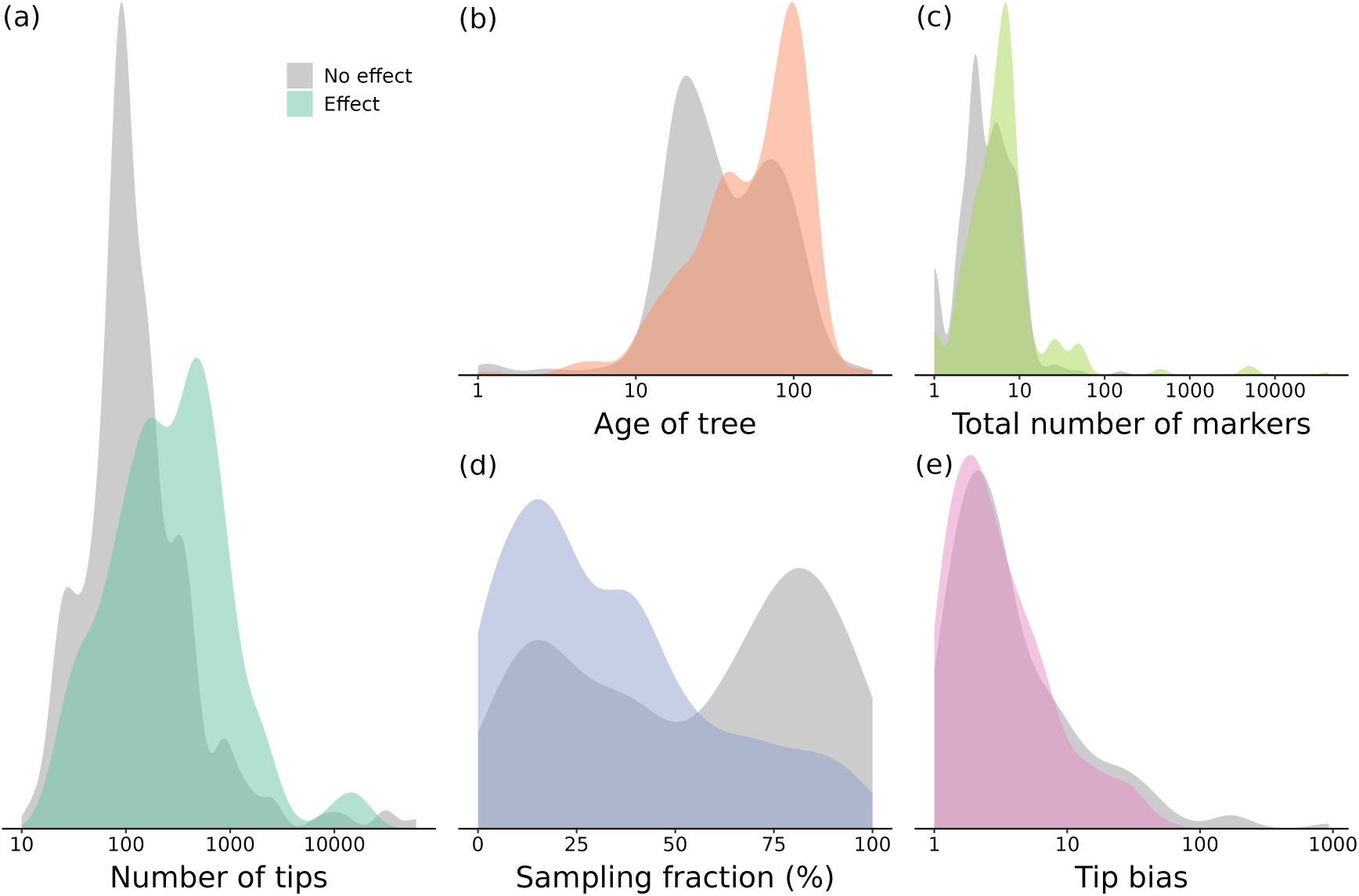
A set of densities depicting the distribution of values for five dataset properties in SSE models that infer trait dependent diversification (coloured densities) and those that do not (grey densities). The dataset properties shown are (a) number of tips in the phylogenetic tree used with the SSE model (data taken from n = 621 models), (b) the age of the tree used with the SSE model (n = 523), (c) the total number of genetic markers (nuclear + plastid + mitochondrial) used to infer the phylogenetic tree used with the SSE model (n = 615), (d) the global sampling fraction (n = 616) and (e) the tip bias, here calculated as the largest tip frequency divided by the smallest (n = 429).

Tree size and root age are closely linked because trees with larger numbers of tips are generally older (Fig. S11). Indeed, we found that trait-dependent diversification was detected more often when trees with an older root age were used (Fig. 4b, S10b). Regardless of their size or age, trees that more accurately represent the true phylogeny of a group will allow us to more reliably estimate its diversification history. We used information on the total number of molecular markers (nuclear + plastid + mitochondrial) as a proxy for tree quality. We found a difference in the distributions indicating that models with trait-dependent outcomes usually had better quality trees than those that did not (Fig. 4c, S10c).

Another issue that has been repeatedly brought up in simulation studies is the potential effect of inflated tip bias (Davis et al., 2013; Maddison et al., 2007). Tip bias increases when there is a higher frequency of one state than the others across the tips of the tree. Upon examining the data used with SSE models we found substantial overlap between densities (Fig. 4e) except for extreme values of tip bias where SSE models tended to find no effect of the trait studied (Fig. S10e). Tip ratio bias recommendations were also made by Davis et al. (2013), who cast doubt on inferences made when the rarest state occurs in less than 10% of the taxa. Prior to 2014, 83% of SSE models (55 of 66) had suitable tip ratios and this figure remained similar (87%, 313 of 360) for the studies that came after.

Global sampling fraction is the proportion of known species that are present in the tree. If the sampling fraction is low it can drastically affect diversification rate estimation (Chang et al., 2020; FitzJohn et al., 2009; Sun et al., 2020). The sampling fraction was explicitly modeled in SSE methods by Fitzjohn et al. (2009), who recommended that the sampling fraction should be at least 25% to adequately capture diversification dynamics. In our data set, sampling fraction ranges from *<*0.1% to complete (100%) sampling. All 10 models published in 2009 and earlier had sampling fractions greater or equal to 25% compared to 60% of 606 models after its publication. This trend (Fig. S12) probably reflects easing of assumptions on complete species sampling, but also indicates that high levels of incomplete sampling are common in recent literature.

Furthermore, we found a striking pattern, showing that those models that used trees in which sampling fraction was low generally yielded trait-dependent diversification, particularly when sampling was less than 40% (Fig. 4d, Fig. S10d). Conversely, high sampling fraction was more often associated with a lack of trait-dependent diversification. Given that the inference of trait-dependent diversification varies with tree size (Fig. 4a), we wondered whether there may also be a relationship between sampling fraction and tree size. However, upon examination we found only a weak, negative trend where trees with more tips had slightly lower sampling fractions (Fig. S13). We then looked at the relationship between sampling fraction and the number of species in the study clade of interest and found a steeper negative relationship (Fig. S14) meaning that the larger the clade of interest is, the less well-sampled it tends to be. Datasets of small clades with low sampling fraction generally dont exist (as they should not be studied) and large clades with high sampling are currently very rare, causing points in the bottom left and top right of figure S14 to be missing. These negative trends remain similar regardless of whether trait-dependent diversification is inferred or not.

### How predictable is the inference of trait-dependent diversification?

Empirical results in angiosperms clearly exhibit strong relationships between various dataset properties and whether trait-dependent diversification is inferred by the SSE model. To assess the importance of the continuous dataset properties together we fit a GAM including the number of tips, root age, the number of markers, the percentage of sampling and the tip bias (Fig. S15). We found that all variables except age of tree were significant when predicting SSE model outcome (*r*^2^ = 0.239, see Table S2 for full details).

If we had comprehensive information about the input data, including the dataset properties investigated above but also information about taxonomy and traits, could we predict whether trait-dependent diversification would be inferred? Using a machine learning approach, extreme gradient boosting (Chen & Guestrin, 2016), we were able to correctly predict, with approximately 72% accuracy (60-80%, Fig. S16), whether SSE models would infer trait-dependent diversification. The most important factors were the information-dense, continuous variables (Fig. 5), further reinforcing earlier observations about their potential influence on SSE model outcomes (Fig. 4, S10, Table S2). Generally, categorical variables related to the trait studied (e.g. fruit morphology), SSE model used (e.g. HiSSE) and order investigated (e.g. Poales) played a smaller but still important role in the model’s predictive ability (Fig. 5).

**Figure 5:**
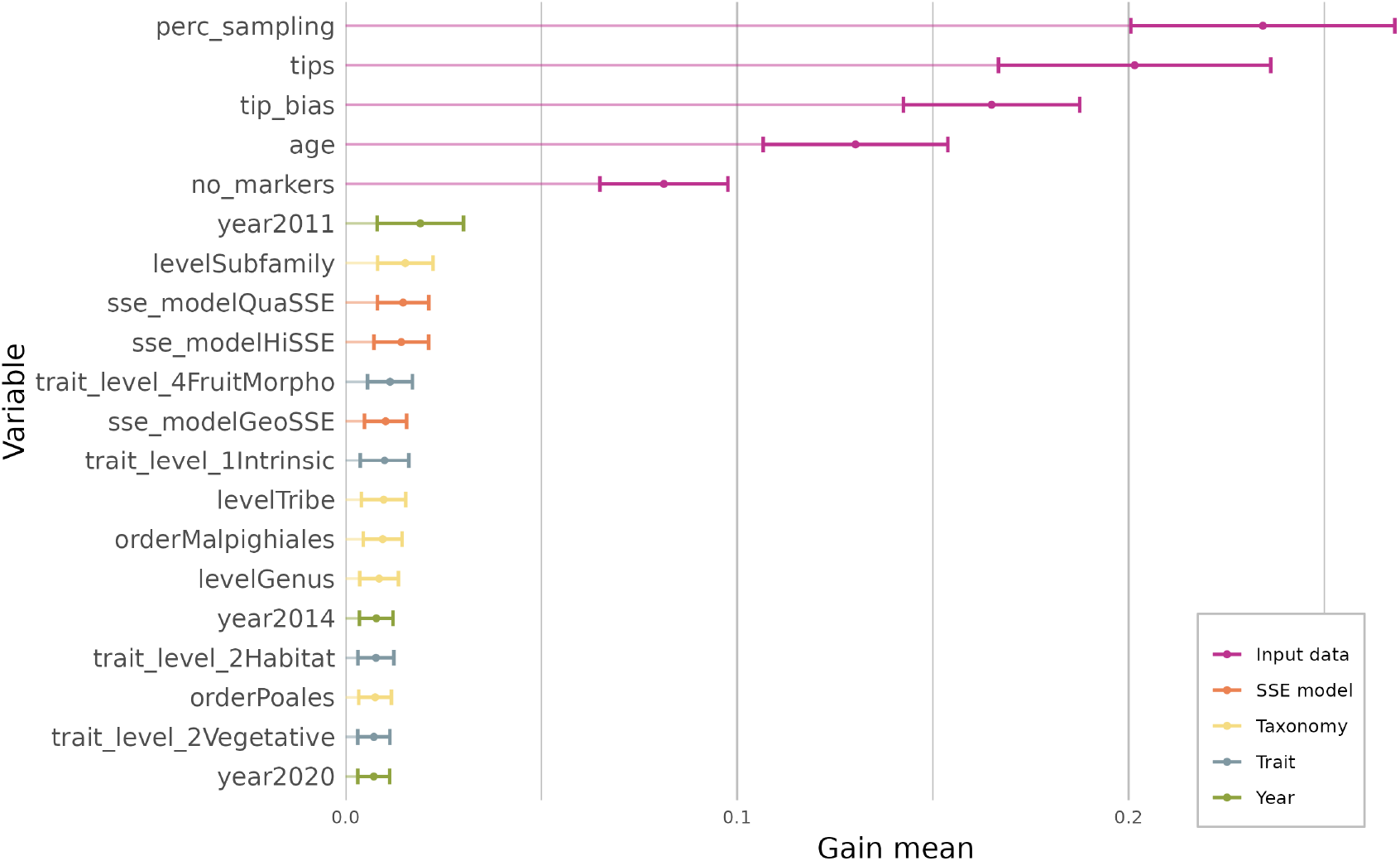
A horizontal barplot showing the relative influence of the 20 most important features included in the xgboost model used to predict the outcomes of SSE models, whether or not trait-dependent diversification is inferred, using input data properties and other characteristics of each study. Points are the mean gain values calculated from the 500 iterations that were run. Error bars represent one standard deviation around the mean. Bars are coloured based on the type of variable they represent. Variable are named using the column headers in the dataset (see appendix S1). For example ‘levelPoales’ indicates that the variable was the category ‘Poales’ from the ‘level’ column.

## Discussion

### No consistent drivers of angiosperm diversification

Previous work has proposed that diversity in different angiosperm groups may have been shaped by various combinations of ecology, traits and environment (Davies et al., 2004; de Queiroz, 2002; Donoghue, 2005; Donoghue & Sanderson, 2015; Hernández-Hernández & Wiens, 2020; Magallón & Castillo, 2009). Indeed, our compilation of the results of 152 studies on trait-dependent diversification in angiosperm clades supports this proposal; that is, the factors driving angiosperm diversification are more complex than a set of universal drivers. When we compared studies investigating the same trait types we found that conclusions generally differed with some indicating that the trait does have an effect on diversification and others concluding there is no effect. We note that the inconsistency observed might reflect real trends in the data, or be due to dataset properties (lack of power, model mis-specification). In the following, we will first discuss the biological conclusions of our study, before considering dataset and model properties.

Our analyses and results centered around how traits (e.g. pollination-related traits) rather than the character states of these traits (e.g. bee vs bird pollination) affect diversification. After grouping traits into several levels of categories (Table S1), we found that some types of traits were more often found to affect angiosperm diversification than others. It may come as no surprise that floral traits, are among the most investigated and influential (Fig. 2b). Indeed, the flower contains the organs needed for sexual reproduction, making it central to the biology, ecology and evolution of angiosperms, and flower characteristics certainly have a large role in determining differences in diversification (Vamosi et al., 2018). In particular, reproductive systems are highly variable in angiosperms (Barrett, 2013) and are again thought to be closely linked to their success (Barrett et al., 1996). Results from SSE models lend some support to this idea - for example, trait-dependent diversification was commonly inferred when mating system traits were investigated (Fig. S5). Even so, we found that in most cases, breeding system (the higher level trait classification including all aspects of mating and sexual systems) often did not yield trait-dependent diversification, due to variability in the effects of sexual systems. Vegetative traits (those related to the growth and non-floral morphology of the plant) and other intrinsic traits including those related to photosynthesis and the genome have received less attention than floral traits (Fig. 2). However, they were more consistently associated with trait-dependent diversification than reproductive traits.

Unfortunately we could say little about which state was advantageous for a given trait because a lack of overlap among states across the 152 studies. Nevertheless we were able to examine how particular states affected diversification in eight traits. Five of these demonstrated how different states of the same trait (e.g. woody and herbaceous species) can increase diversification in different groups (Fig. S5) meaning that only three showed consistent patterns where one state was associated with elevated diversification rates (epiphytism, non-C3 photosynthesis and self-incompatibility). However it is difficult to say whether these have truly consistent effects on diversification as they have only been investigated a handful of times in a relatively small proportion of angiosperm species (e.g. self-incompatibility has been tested in Solanacaeae and Onagraceae only).

Despite some general patterns in those traits that are more often influencing angiosperm diversification, the overarching trend is that the effect of a trait on diversification is clade-dependent. Therefore, the main question remains open: what drives differences in diversification among angiosperms? The fact that a definitive answer has yet to be found suggests that it’s the complex interplay between trait evolution, biotic interactions and geography that matters. Indeed, geography (range size, biome) has been identified many times as an important factor (Hernández-Hernández & Wiens, 2020; Vamosi et al., 2018), but it is unclear whether this is a cause or a consequence of differences in diversification. Others have suggested that it is not the presence or absence of a trait that determines the evolutionary success of a clade, but rather the capacity to change (Onstein, 2019; Ricklefs & Renner, 1994). This could partly explain the inconsistency of the inferences, but again, trait diversity could be both a cause and a consequence of species richness. Furthermore, the choice of clades and traits, as well as the quality of the input data, also influence whether or not differences in diversification are detected, and therefore our conclusions.

### The importance of evolutionary scale and context

Users of models of trait-based diversification face an important challenge - choosing the context in which to conduct analyses. In the simplest scenario, where a trait only evolved once in the study clade, its effect on diversification cannot be tested (Maddison & FitzJohn, 2015) and thus this type of context should be avoided. At the intermediate scale a trait may have evolved multiple times in closely-related clades but their evolutionary context (i.e. species’ genomes, morphology, ecology, or external environments) is much more similar than distantly-related ones. So, associations between states and rates cannot be interpreted as a general pattern in this limited phylogenetic scope. Broadening the scope of the analysis, by way of either a larger tree, or multiple trees in a meta-analytic framework (Sabath et al., 2016) can help to reveal these general patterns but leads to different challenges.

At larger phylogenetic scales, trees with many taxa are more likely to contain a range of branching patterns where lineage accumulation is faster in some parts of the tree than in others. In older clades there has been more time for macroevolutionary processes to have an impact on the trees we infer and the traits we observe today. As we observed, larger, older trees more commonly yield trait-dependent diversification (Fig. 4). However, their heterogeneity (due to e.g. molecular clock rate variation; Shafir et al., 2020) would also make them more susceptible to false-positive errors that could over-inflate the number of times trait-dependent diversification is detected (Rabosky & Goldberg, 2015). Indeed, one of the major criticisms of early SSE models was the propensity to infer false positives due to model inadequacy: the models were based on the assumption that only the trait of interest would influence diversification, so any kind of heterogeneity would lead to the rejection of the null hypothesis (Maddison & FitzJohn, 2015; Rabosky & Goldberg, 2015). This could explain the inconsistency of the effects of traits across clades - it may be that false positives caused by lineage-specific factors correlated with the shared focal trait are driving the disparate patterns. Models with hidden states go some way towards alleviating this issue as they can account for lineage-specific factors. When only considering results from models with hidden states we found that the proportion of trait-dependent diversification changed substantially for some traits, though inconsistency was still common (Figs. 2, S8). As more studies with hidden states are conducted, we will find out whether these trends are general. While hidden states models certainly are an improvement, they assume that these states are categorical and have constant transition rates, which very likely doesn’t capture all sources of heterogeneity. They cannot handle all cases of possibly misleading inferences, e.g. when the effect of a trait that evolved multiple times is driven by one clade where it strongly influences diversification while leaving it unchanged in other clades (Beaulieu & O’Meara, 2016; Maddison & FitzJohn, 2015). Furthermore, there has yet to be a study that thoroughly assesses the model adequacy of HiSSE, as has been done for BiSSE (Rabosky & Goldberg, 2015).

### Best practices for SSE model use and result reporting

Though a number of recommendations have been made for accurate inference with SSE models, most empirical studies do not meet them. When using the strict thresholds suggested in the literature (25% taxon sampling, 300 tips and minor tip state frequency of 10%; Davis et al., 2013; FitzJohn et al., 2009) we find that just 20 of 152 studies contain models that meet all criteria. The apparent relationship between sampling fraction and inference of trait-dependent diversification (Fig 4) should invite us to be cautious about studies using low sampling fractions, as it has been shown that better sampled trees yield more accurate estimates of diversification rates (Chang et al., 2020; FitzJohn et al., 2009). However, publication bias may also be playing a role. If no trait-dependent diversification is detected in a poorly sampled clade this may be attributed to a lack of power that ultimately prevents publication, thereby inflating the number of studies with low sampling that detect trait-dependent diversification. To clarify these observations, simulation studies should be undertaken to investigate the influence of sampling fraction together with model inadequacy on SSE model inference.

In most studies, some of the information we consider crucial for the interpretation of the results was lacking, or it was difficult to access. Collecting data for many properties (e.g. samples per state) required us to count from figures or extract statistics from archived raw data, which were not always freely available. For example, we were unable to extract and use the number of independent origins of each character state. Robust estimates of associations between traits and diversification rates necessitate multiple independent origins (but not too many (Rabosky & Goldberg, 2015)) and corresponding rate changes (FitzJohn et al., 2009), so an idea of this value per study, inferred using ancestral state reconstructions, would be useful for interpretation of the robustness of results. This could be done by combining stochastic mapping of traits with an SSE model (Freyman & Höhna, 2019), though this is generally not available for SSE approaches. Likewise, diversification and transition rates were often not reported in an easily-accessible and standardized manner, or in some cases, not at all. These should be reported, and if possible, with confidence metrics around rate estimates e.g. Bayesian credible intervals.

Louca & Pennell (2020a) recently pointed out how diversification rate estimation can be susceptible to issues of unidentifiability. Though SSE models are not directly implicated (Helmstetter et al., 2021), one potential way to help ‘future proof’ analyses from unidentifiability caused by overfitting would be to avoid reporting and assessing speciation and extinction rates separately, focusing instead on compound parameters such as net diversification rate (*λ* − *μ*), turnover rate (*λ* + *μ*) and extinction fraction (*μ/λ*) that are typically used in more recent SSE models (e.g. HiSSE). To encourage standardized result reporting we propose an initial set of characteristics that should be made available in all future studies using SSE models (Supplementary Data 1).

Given that evolutionary context appears to be important for understanding trait-dependent diversification, how to best choose a trait and clade to study? Trait choice can be helped by preliminary knowledge of the phylogenetic tree and ancestral state reconstruction, which could be used to ensure that the derived state(s) arose multiple times and that the ratio among different states is not extreme (*<*10:1). In terms of choosing a clade, it is first important to adhere, as best as possible, to the recommendations for using SSE models e.g. avoid clades much smaller than 300 taxa and focus on those that are well sampled (*>*25%). If recommendations cannot be followed, because of natural limitations in clade size, for example, these should be stated clearly as caveats.

Working at a much larger scale, e.g. angiosperm-level analyses, is certainly appealing but creates a range of issues related to confounding factors that current models will find difficult to disentangle. To better learn about the factors that influence angiosperm diversity we therefore suggest studies focus on multiple intermediate-sized clades i.e. large genera, families or tractable orders. However, if these clades are well-sampled they would approach the limit of our current computational feasibility (but see Louca and Pennell, 2020b). Working with many smaller clades may therefore be more feasible in the near future and also yield important insights via the comparison of diversification patterns among many different groups (e.g. Sabath et al., 2016), which we think is an acceptable tradeoff for reduced power in standalone analyses. Examining the effect of the same trait in multiple clades would allow researchers to account for the unique and shared aspects of their biology (e.g. through the use of hidden states or trait combinations), and then to combine results (Rabosky & Goldberg, 2015) to uncover general patterns.

### Knowledge gaps and future avenues

Our review allowed us to identify groups that are understudied and therefore good focal points for future research to gain a more well-rounded picture of angiosperm macroevolutionary dynamics. One of the most obvious is Asteraceae, species-rich yet subject to relatively few trait-based diversification studies (Fig. S4), or Alismatales, an order that has more than 4,500 species (Fig. S3) but just a single study on their trait-based diversification (Canal et al., 2019). In addition, some families with more than 1,000 species, such as Phyllanthaceae or Orobanchaceae have yet to be studied in this way.

High-quality phylogenetic trees are not the only ingredient for SSE studies; trait data also need to be available. We highlight traits related to lifespan, dispersal and symbiosis as ripe avenues for future work that have potential to unearth important patterns in trait-dependent diversification. However, apart from a few traits such as geographical range or climatic preferences, gathering high-quality data for large numbers of species is a time-consuming activity. We encourage the integration of trait data generated from SSE studies (and others) into large, global trait databases such as eFLOWER (Sauquet et al., 2017), TRY (Kattge et al., 2020) or more focused databases (e.g. AusTraits (Falster et al., 2021)). These will act as important resources as researchers consider several traits in tandem when testing for context-dependent effects of traits, or when disentangling the traits hiding in the hidden-state approaches. Most importantly, studies should be conducted on traits where clear hypotheses can be generated about their effect on diversification in the chosen study clade.

SSE methods are statistical tools that are aimed to uncover correlations, and cannot themselves discover causal relationships. By definition, macroevolutionary models try to capture the result of many aggregated small-scale processes in a few high-level parameters. Speciation is an instantaneous split of one branch into two in most macroevolutionary models, although in reality there might be a wide range of different dynamics depending on environmental heterogeneity, biotic interactions, and intrinsic traits (e.g. breeding systems, genomic incompatibilities) (Coyne & Orr, 2004). Thus, if a trait is predicted to affect speciation and extinction, high-quality inferences of diversification rates, for which our synthesis provides some guidelines, should be able to detect a signal. However, this signal is only a piece of the puzzle, as it is through various ecological and genetic processes that can also be put to the test. For example, Park et al. (2018) compared sister species with contrasted mating systems (selfing vs. outcrossing) and showed that niche breadth tended to decline over time in selfing lineages, in agreement with the dead-end scenario proposed for this trait and detected in macroevolutionary analyses (Goldberg & Igić, 2012; Höhna et al., 2019). Additionally, we can identify traits that have an effect on ecological and genetic mechanisms that control speciation and extinction, such as traits affecting coexistence and niche partitioning (Adler et al., 2013) (e.g. specific leaf area or seed mass), genetic differentiation between populations or species (Gamba & Muchhala, 2020) (e.g. pollination mode, mating system, growth form) or those associated with commonness and rarity (Murray et al., 2002) (e.g. seed production). Such traits come with *a priori* hypotheses and could be ideal candidates for macroevolutionary studies exploring their effect on diversification.

## Conclusions

When bringing together the last 12 years of study on trait-dependent diversification in angiosperms, it is the inconsistent effects of traits that stand out, rather than the importance of a particular set of universal drivers. This highlights the important role the evolutionary context of a clade plays in determining how a particular trait affects diversification. Furthermore, the nature of the data itself, relating to factors such as how well-sampled or large a clade is, was shown to have substantial influence on SSE model results. The guidelines we set out in this review will help to improve how we use trait-dependent models and our template for reporting results will facilitate future synthesis as SSE models continue to be used and developed. We have only touched the surface of what we can learn about trait-dependent diversification in angiosperms. Will results from novel studies change the trends we observe here? Given the production of new datasets that meet recommendations for robust inference, future methodological developments enabling studies at wider scopes and the potential for new discoveries in understudied traits and clades, we think it is certainly possible. Though our study focused on angiosperms the conclusions we draw about consistency, context dependence and SSE model use will apply to studies of trait-dependent diversification across the entire tree of life.

## Acknowledgements

We thank the DiveRS group for the ideas and discussions that led to this manuscript. We thank Nicolas Casajus and Aurore Receveur for advice on data analyses.

## Supplementary Material

Code and data associated with this manuscript is available from http://github.com/ajhelmstetter/sseReview. PapieRmache can be found at http://github.com/ajhelmstetter/papieRmache.

