## supplementary materials for "Trait-dependent diversification in angiosperms: patterns, models and data"

[1]*Fondation pour la Recherche sur la Biodiversité - Centre for the Synthesis and Analysis of Biodiversity, 34000 Montpellier, France.* , [2]*School of Life Sciences, University of Hawaii Manoa, Honolulu, HI, 96822, USA.*, [3]*National Herbarium of New South Wales, Royal Botanic Gardens and Domain Trust, Sydney, New South Wales, 2000, Australia*, [4]*Evolution and Ecology Research Centre, School of Biological, Earth and Environmental Sciences, University of New South Wales, Sydney, Australia.*, [5]*Department of Zoology, 6270 University Blvd., University of British Columbia, Vancouver BC, V6T 1Z4, Canada.*, [6]*Area of Biodiversity and Conservation, Universidad Rey Juan Carlos, 28933 Móstoles (Madrid), Spain.*, [7]*Department of Biological and Environmental Sciences, University of Stirling, Stirling, FK9 4LA, United Kingdom.*, [8]*Department of Botany and Biodiversity Research, University of Vienna, Rennweg 14, 1030 Vienna, Austria.*, [9]*Department of Organismal Biology, University of Uppsala, Uppsala, 75236, Sweden.*, [10]*Department of Botany and Zoology, University of Stellenbosch, Private Bag X1, Matieland 7602, South Africa.*, [11]*Natural History Museum, University of Oslo, 0318 Oslo, Norway.*, [12]*CNRS, Ecosystèmes Biodiversité Evolution (Université de Rennes), 35000 Rennes, France.*, [13]*Université de Lyon, Université Lyon 1, CNRS, Laboratoire de Biométrie et Biologie Evolutive UMR 5558, F-69622 Villeurbanne, France.*

### Contents


1 Appendix S1: Dataset characteristics 3

2 Supplementary figures 5

3 Supplementary tables 21

### 1 Appendix S1: Dataset characteristics

Here we give details of all of the different dataset characteristics collected from studies that used SSE models to investigate diversification in angiosperm groups. These characteristics correspond to the column headers in our synthesised dataset as well as the template for reporting SSE results (Supplementary Data 1).

**entrant** The name of the person who entered the data into the spreadsheet.

**authors_contacted** Indicates whether authors of the study were contacted for clarifications.

**author_feedback** Indicates whether a response was recieved from contacted authors.

**study** This is the unique name given to the study, this was taken from the text file that the PDF version of the article was converted to. It is in the format "AuthorYearTitle.txt".

**year** The year in which the study was published. Only peer-reviewed, published studies were included (no preprints).

**sse_model** The name of the state-dependent speciation and extinction (SSE) model used. There may be multiple models of the same (or different) types per study. See Fig. 1 for the range of SSE models considered.

**model_no** A numeric identifier given to each model in each study. This number is repeated for the number of states included in each model. If a study reports results from a BiSSE model and MuSSE model with three states, numeric identifiers would be as follows: "1,1,2,2,2".

**order** The angiosperm order that the study group belongs to. If the scope of the study includes multiple orders (e.g. a study at the angiosperm level) then "multiple" is used.

**trait_level_1-trait_level_6** Character states are classified into trait types at different levels, with level 1 being the most broad, and level 6 being the most specific. This classification follows the trait ontology, which can be found in Table S1.

**character_state** The character states used in a given model. There may be one or more character states per model. GeoSSE and GeoHiSSE models are specifically designed to assess diversification differences among geographic regions and here we only consider states representing the geographic regions, omitting the special state "widespread" used for taxa that are present in both regions.

**putative_ancestral_state** A binary column that indicates the character state which is supposed to be ancestral (1) in the analysis. We use "putative" because in many studies the ancestral state was not explicitly reported. In studies that assumed an ancestral state, we treated this as the ancestral state. In studies that performed ancestral state reconstruction and reported results, we chose the state that was most likely at the root of the tree used with SSE model. In studies that did not report either of these, we searched the text for statements related to which trait was ancestral or examined the distribution of tip states to identify which state is putatively ancestral. Therefore in some cases this characteristic is somewhat subjective and should not be considered as accurate evidence for the ancestral state of the trait, but instead a hypothesis about what the ancestral state is likely to be.

**clade** The name of the clade that the SSE model was run on.

**level** The taxonomic level of the clade that the SSE model was run on (e.g. genus, tribe or order).

**order, family** The family and order the study clades belong to. If larger than family or order the category "multiple" was used.

**div_inc** A binary column indicating when net diversification rate was higher (1) for a character state. We initially wanted to include only significant results here, but in many cases significance was not reported, so we considered all cases in which net diversification was reported to be higher in one trait than the other as trait-dependent diversification. In multi-state models we classified the state with the lowest net diversification rate rate as 0 and other states as 1,2,3... etc depending on their net diversification rate. If two states are explained as having comparable rates in the text they are assigned the same number.

**sp_inc** A binary column indicating when speciation rate was higher (1) for a character state.

**ext_inc** A binary column indicating when extinction rate was higher (1) for a character state.

**no_markers** The total number of nuclear, plastid and mitochondrial markers used to build the phylogenetic tree that was used with the SSE model. Equal weight was given to each marker type in this column (even though plastid markers, for example, are not entirely independent).

**no_plastid** The total number of plastid markers used to build the phylogenetic tree that was used with the SSE model.

**no_nuclear** The total number of nuclear markers used to build the phylogenetic tree that was used with the SSE model.

**no_mito** The total number of mitochondrial markers used to build the phylogenetic tree that was used with the SSE model.

**age** The age of the root of the phylogenetic tree in million years. As with tips, we tried to get the age of the tree that was used with the SSE model if this differed from the age of the tree reported in the main text of the study. However, this was not always possible. In cases where ages were not reported/data was not available we attempted to estimate ages from figures.

**age_inferred** In some cases the phylogenetic tree was not time-calibrated (e.g. the root of the tree was set to a fixed age of 1). This binary column indicates whether age was inferred (1) or not (0).

**tips** The number of tips used with in the SSE model. In some cases only the number of tips in the tree was reported. We tried to remove outgroups/those taxa included in the phylogenetic tree bit not included in the SSE model where possible, but this was not always evident.

**perc_sampling** The global sampling fraction for taxa used in the SSE model. In cases where this was not reported we acquired estimates for the number of species in the clade of interest from http://www.theplantlist.org/ and used this to calculate a sampling fraction.

**sampling_per_state** The sampling fraction per state, as reported in the study. If this was not reported we did not try to calculate it based on other sources of information as it requires specialist knowledge about which species possess each character state.

**samples_per_state** The number of tips that belong to each character state in an SSE model. In cases where this was not reported we counted tip states from figures.

**transition_direction** The direction of the transition rate (e.g. 0 to 1).

**transition_rate** The transition rate between the states indicated in transition direction (lineages per million years).

**div_rate** The net diversification rate (lineages per million years). If hidden states were included in the model this was calculated as the mean across hidden states (e.g. $(r_1A + r_1B/2)$).

#### 2 Supplementary figures

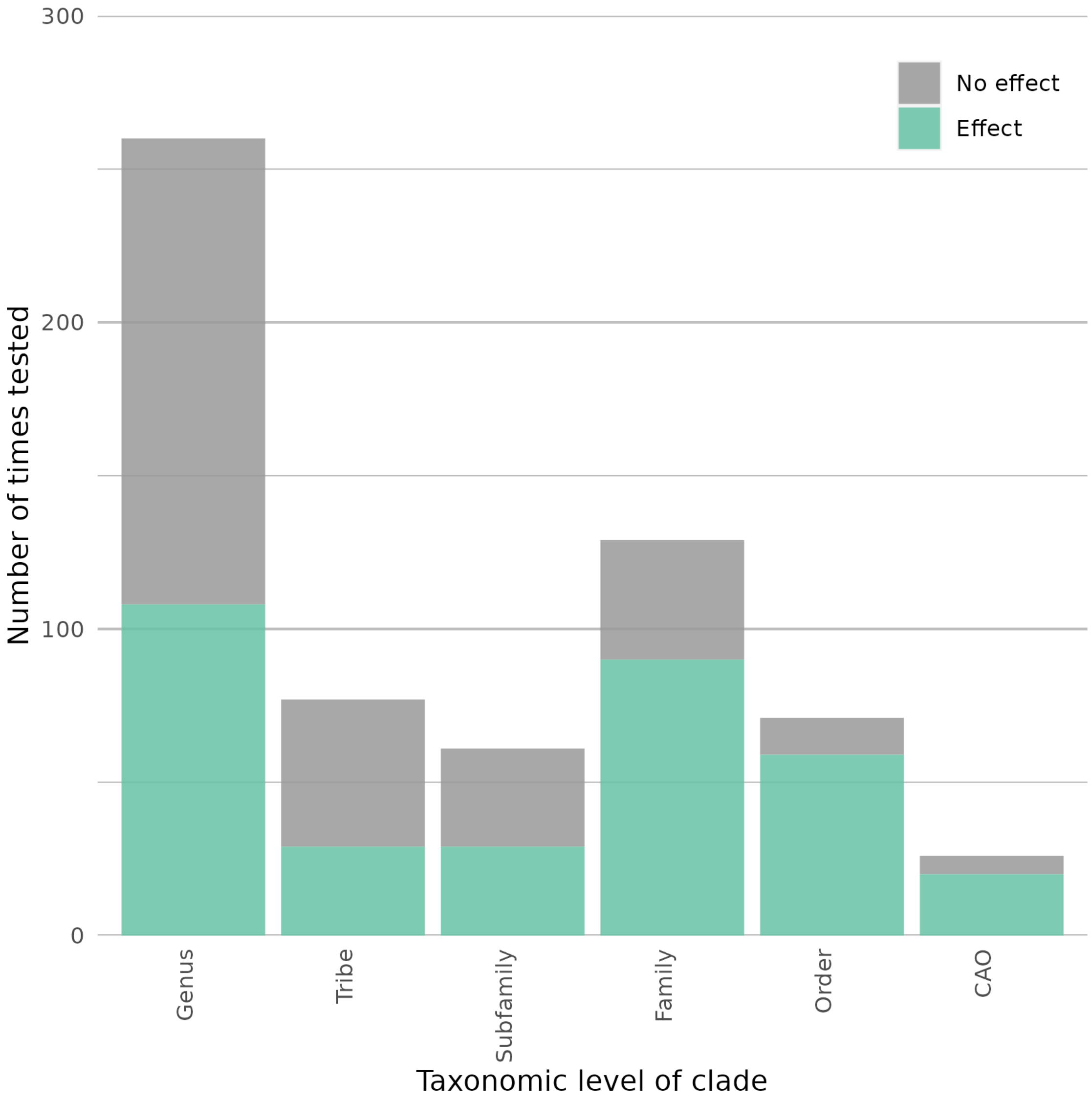


Figure S1: Stacked barplots showing the number of models run per different taxonomic levels. The smallest level is genus, increasing in scope until clades above order (abbreviated as CAO). Note that the category tribe includes subtribes. Bars are coloured based on whether trait-dependent diversification was inferred in the model.

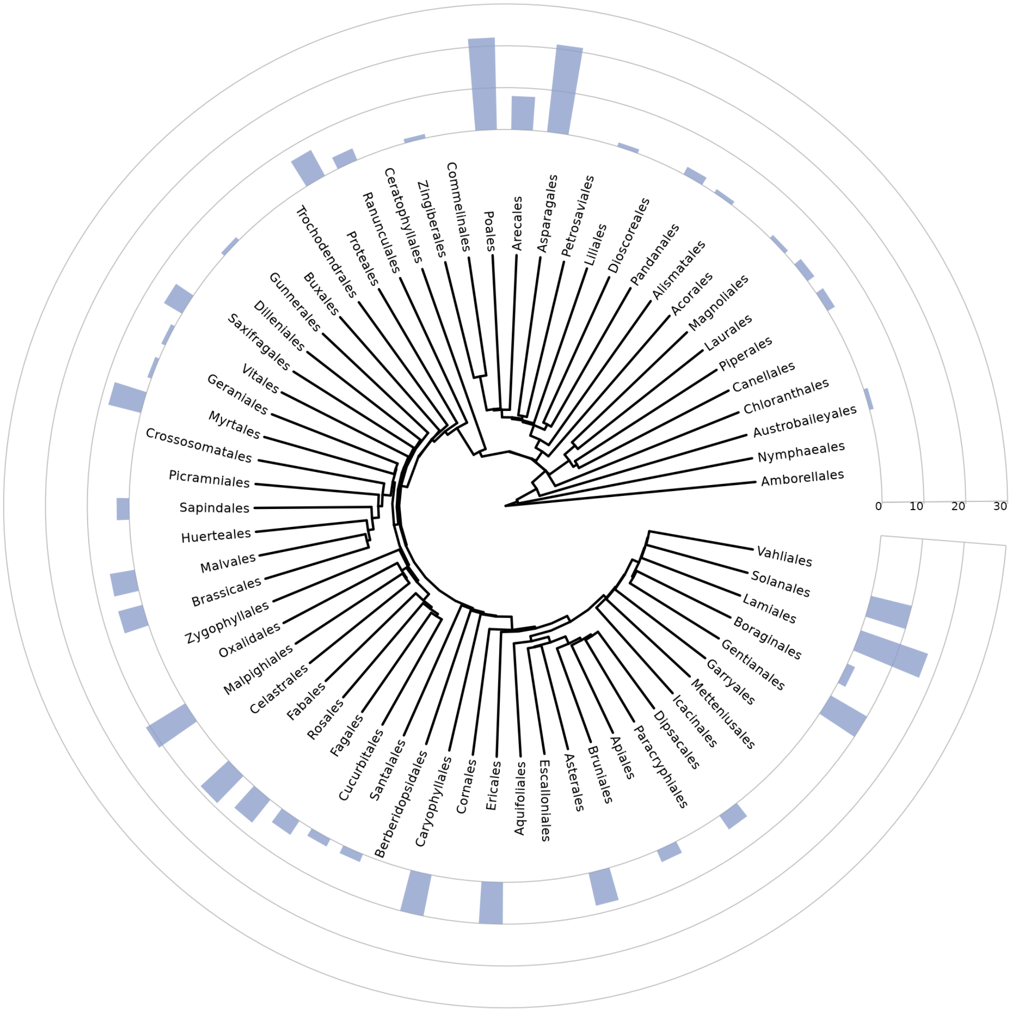


Figure S2: A phylogenetic tree of angiosperm orders taken from Li et al. 2019 Li et al., 2019 annotated with bars representing the number of studies using SSE models that focused on each order.

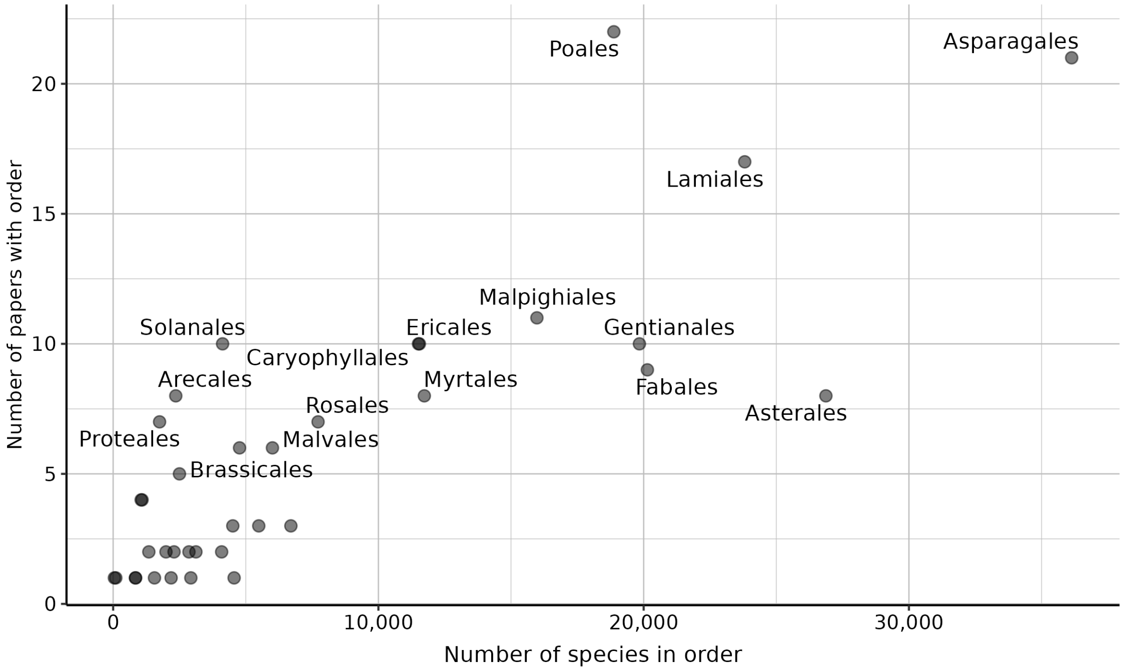


Figure S3: A scatterplot showing the relationship between the number of species in each order (taken from http://www.mobot.org/) and the number of papers that studied that order. Only those orders in our dataset are included. Text labels have been added to the points corresponding to the most commonly studied orders in our dataset.

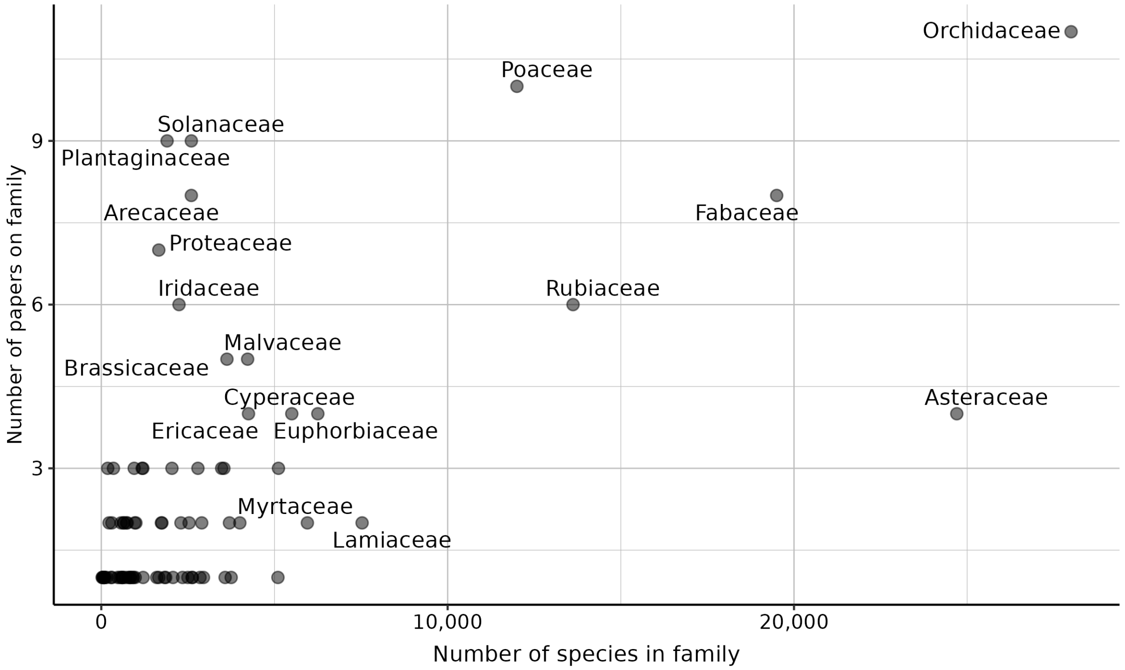


Figure S4: A scatterplot showing the relationship between the number of species in each family (taken from Christenhusz and Byng, 2016) and the number of papers that studied that family. Only those orders in our dataset are included. Text labels have been added to the points corresponding to the most commonly studied orders in our dataset.

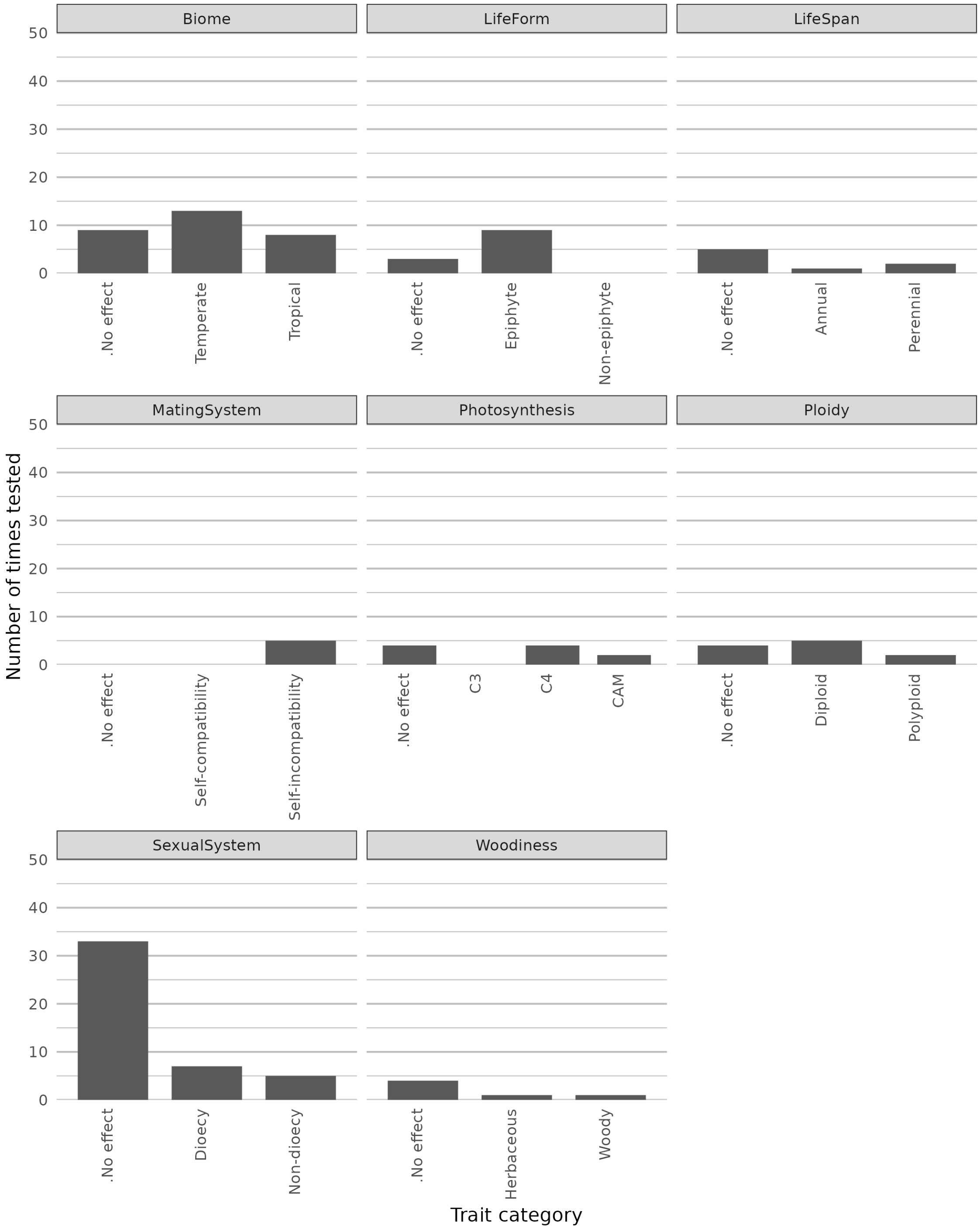


Figure S5: Stacked barplots for eight different character state groups. For each character state there is a bar representing the number of models that state was found to increase diversification rate. If there was no effect of either state in a model, this was counted in the no effect bar. Though the photosynthesis category has three states, these were only tested with binary-state models (e.g. C3 vs C4 or CAM).

.

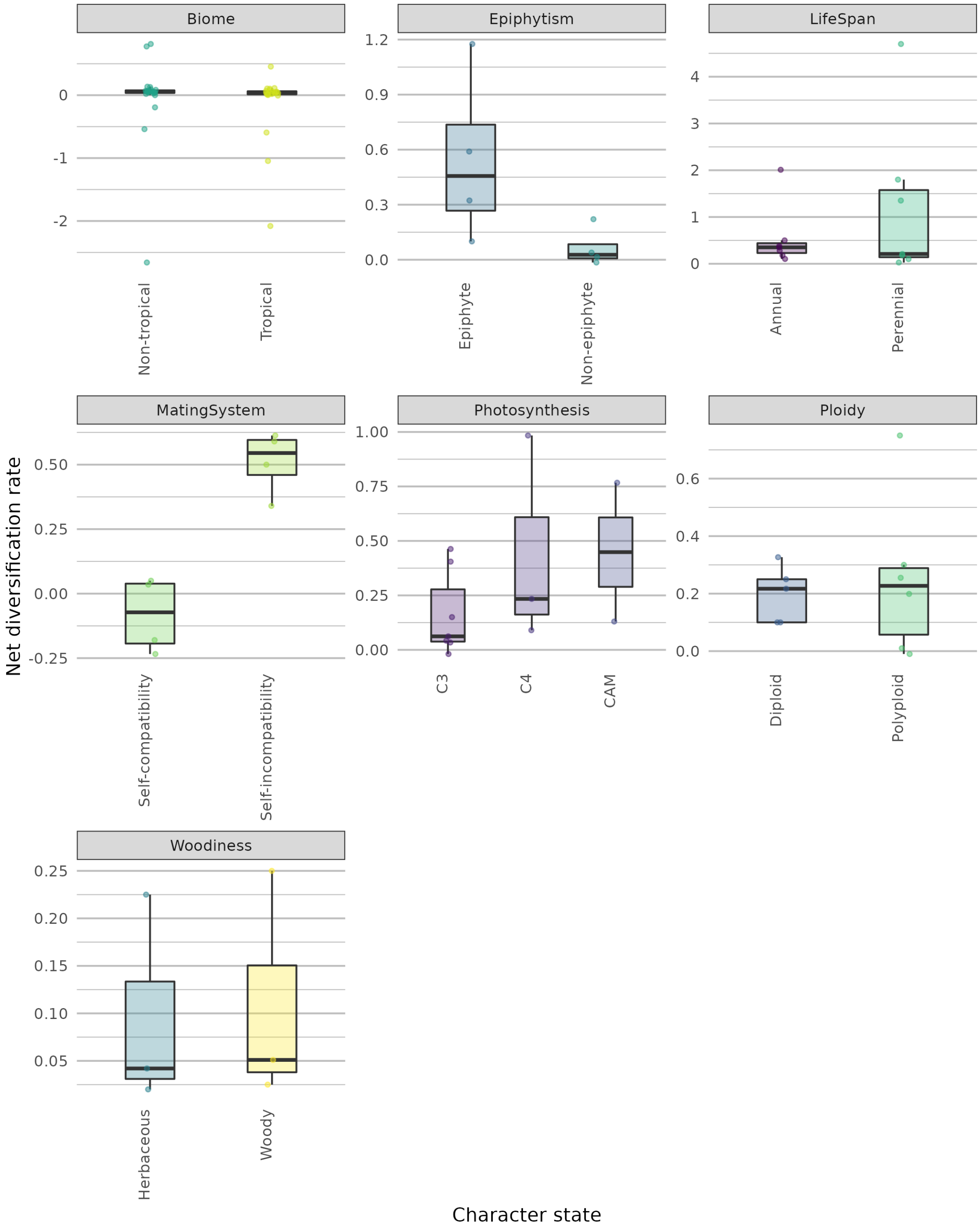


Figure S6: Boxplots showing the net diversification rates for seven traits and their associated states. Jittered points overlain on the boxplots indicate the mean net diversification rate values recovered from each model. The "LifeForm" trait level 6 category was separated into "Epiphytism" and "Woodiness" for presentation purposes.

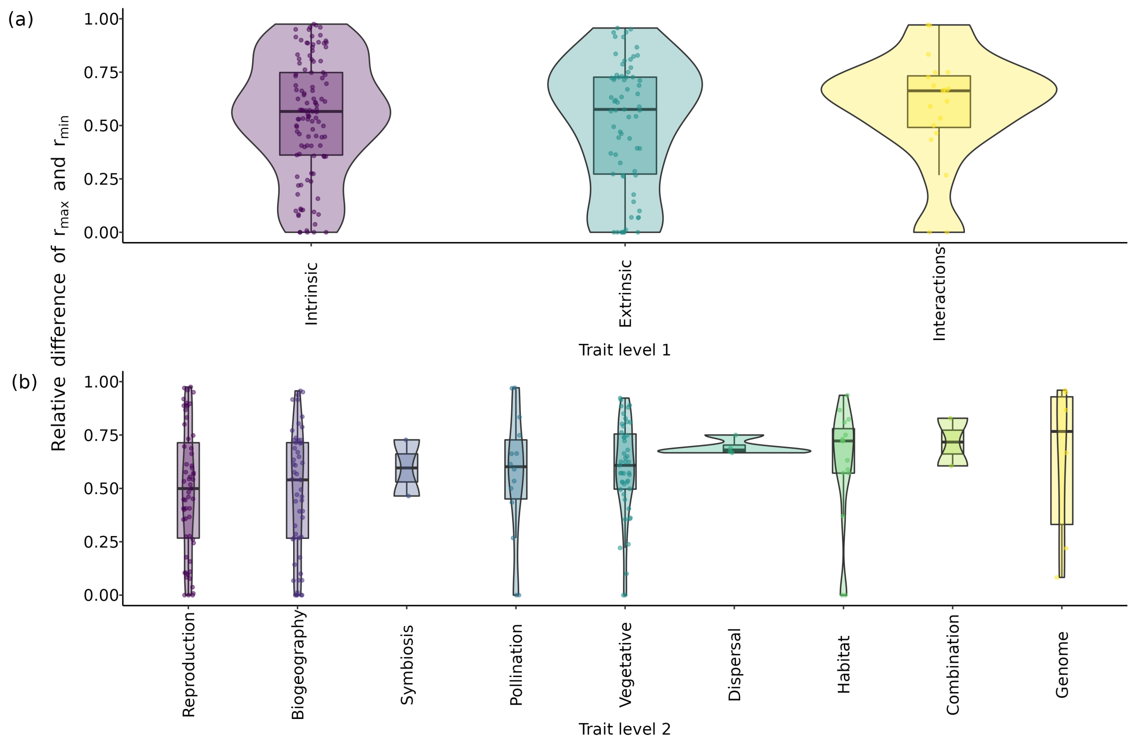


Figure S7: Violin plots showing relative difference between minimum and maximum net diversification rates for models belonging to (a) trait level 1 categories and (b) trait level 2 categories. We calculated the ratio using the formula $(r_{max} - r_{min})/r_{max}$. Only those models in which all rates were positive were used (n = 202). Jittered points overlain on the violin plots indicate the ratio values recovered from each model. For (a) trait level 1 we found rates were generally similar with some indication that interaction traits may have a greater effect on diversification rates than intrinsic or extrinsic traits, but this was not significant (Kruskal-Wallis chi-squared = 0.993, df = 2, p-value = 0.609). For (b) trait level 2, traits related to the genome had the largest relative differences but again categories were not significantly different (Kruskal-Wallis chi-squared = 9.118, df = 8, p-value = 0.333).

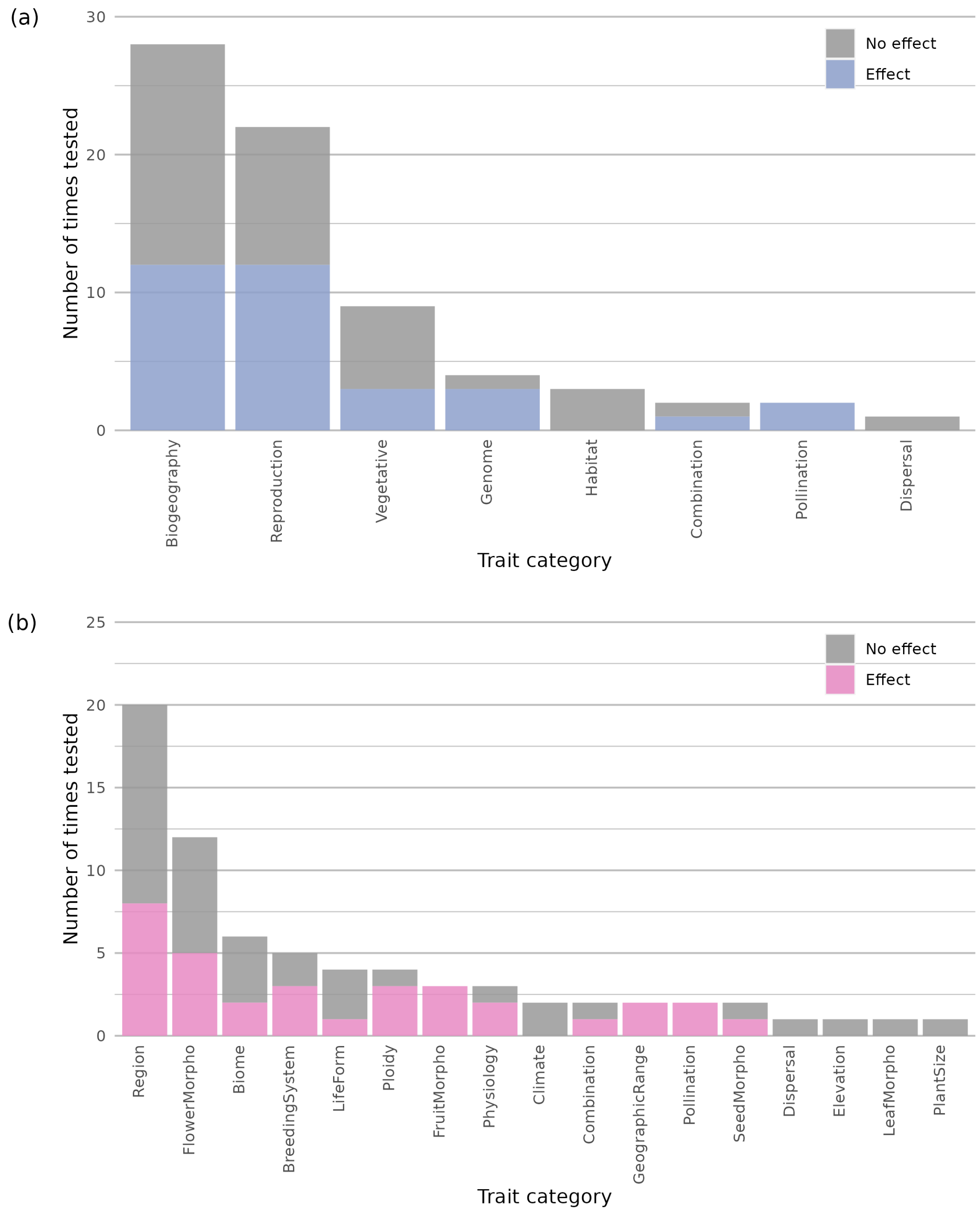


Figure S8: Stacked barplots showing how often particularly trait types were tested, for models with hidden states only. Bars are coloured to depict how often trait-dependent diversification was detected per trait type. If multiple state-dependent speciation and extinction (SSE) models were used in a single study they were considered cumulatively. Two plots are shown, (a) one with broad level 2 trait categories and (b) one with more narrow level 4 categories. An ontology depicting how different trait classification levels are connected can be found in Table S1 and a similar figure including information from all SSE models in our dataset can be found in Figure 2 in the main text.

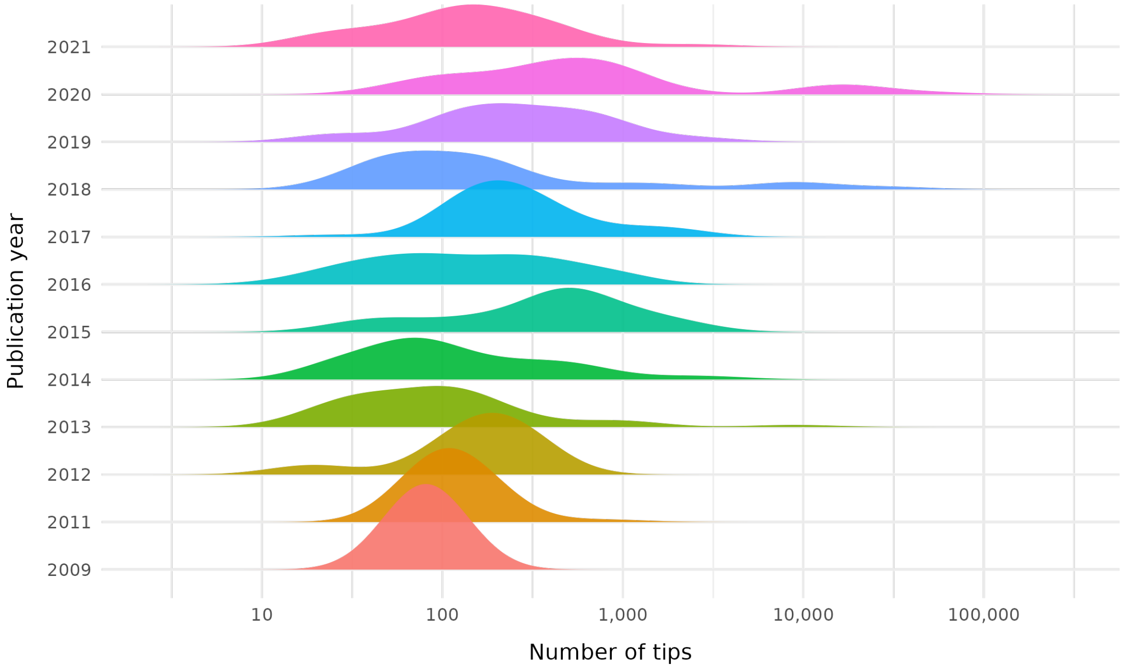


Figure S9: A ridgeplot showing how the number of tips on trees used with SSE models have changed over time. Each ridge displays a density plot corresponding to a single publication year (2009-2021) with the most recent year at the top of the plot. The x-axis is on a log scale. There was not enough data from 2010 to calculate a density so this year was removed from the plot.

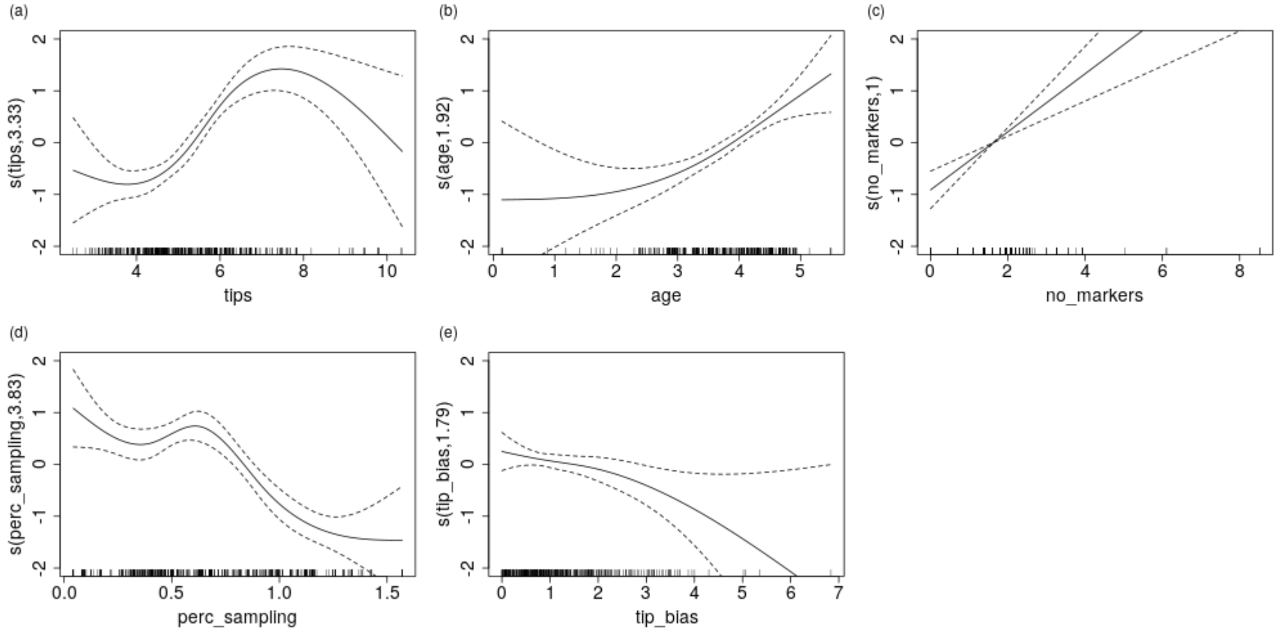


Figure S10: Panels (a-e) show the relationships inferred with generalized additive models (GAM) between each continuous dataset property in Figure 4 and SSE model outcome. Values that are positive on the y-axis indicate that the dataset property at this x-axis value tends to be associated with trait-dependent diversification. Negative y-axis values indicate when dataset properties are associated with no effect. Cubic regression splines were fitted to each property independently.

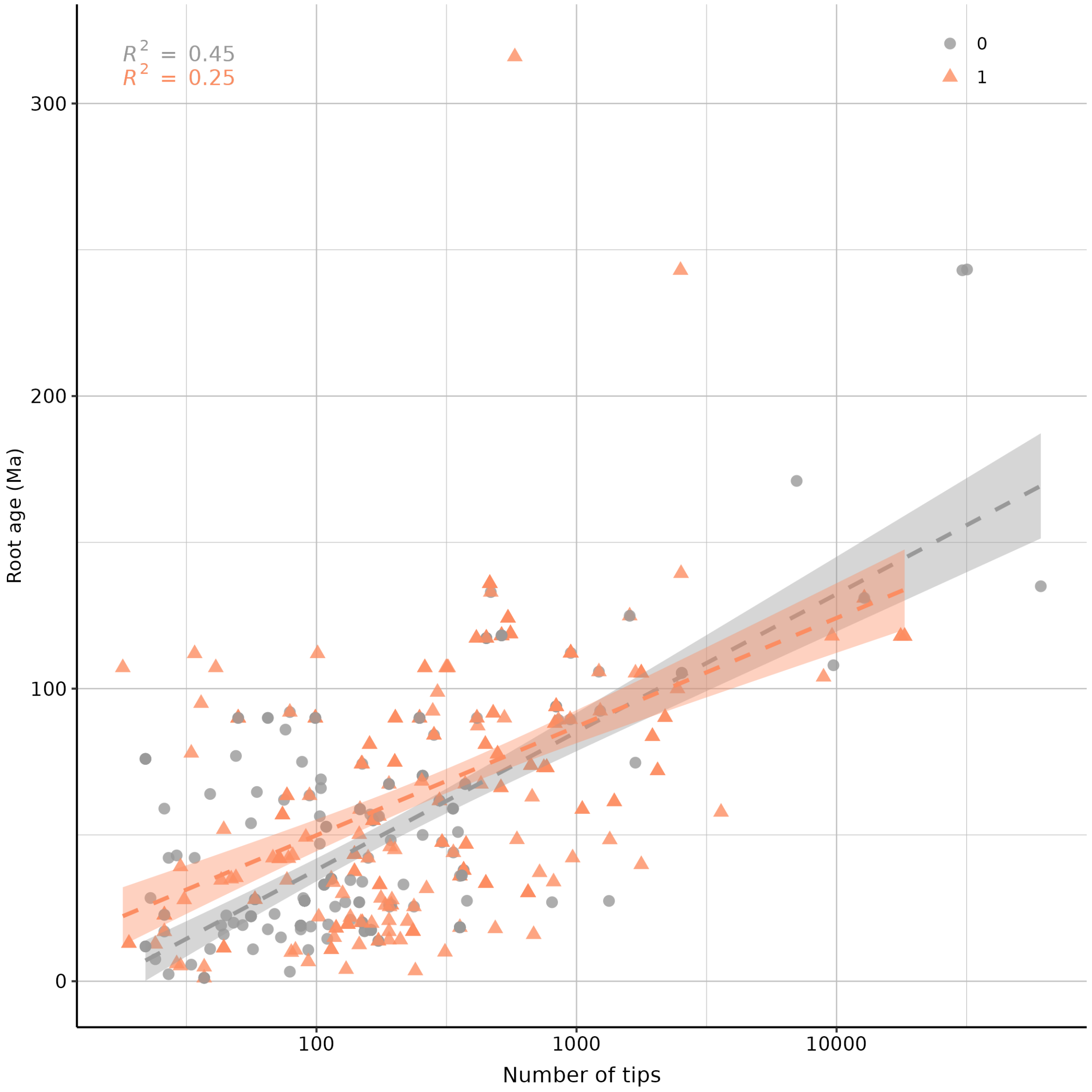


Figure S11: A scatterplot showing the relationship between the root age of the trees used with SSE models in our dataset and the number of tips in the trees. Points are coloured based on whether trait-dependent diversification was inferred (coloured) or not (grey) when the associated model was run. Lines were fitted using linear models to these two groups with 95% confidence intervals estimated.

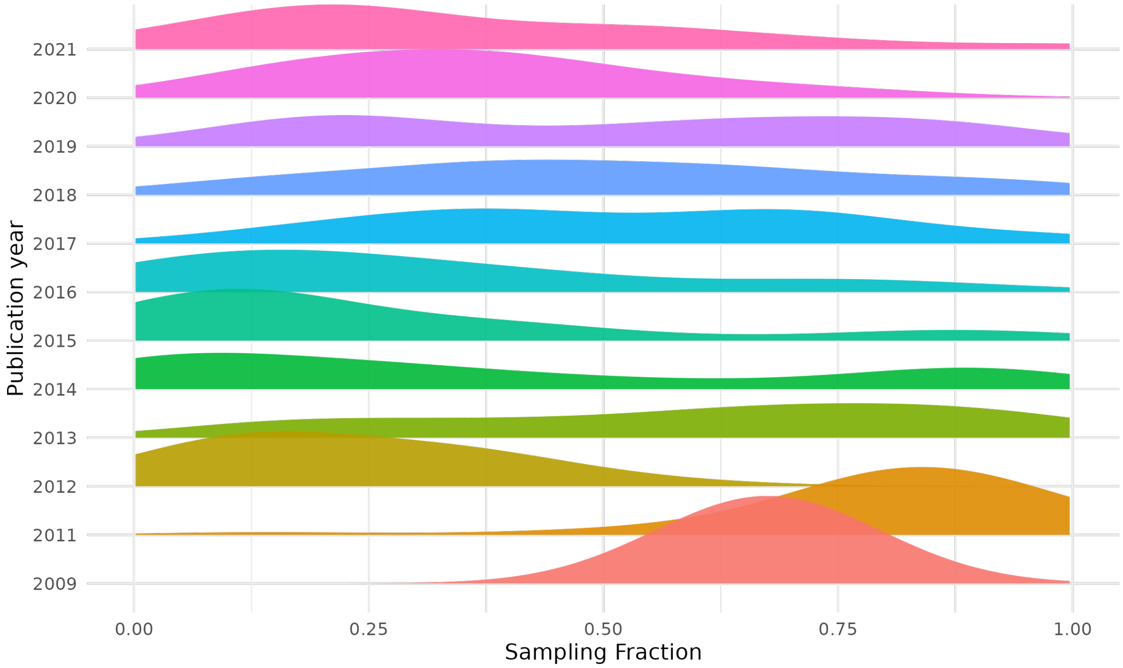


Figure S12: A ridgeplot showing how sampling fractions of trees used in SSE models have changed over time. Each ridge displays a density plot corresponding to a single publication year (2009-2021) with the most recent year at the top of the plot. The x-axis is on a log scale. There was not enough data from 2010 to calculate a density so this year was removed from the plot.

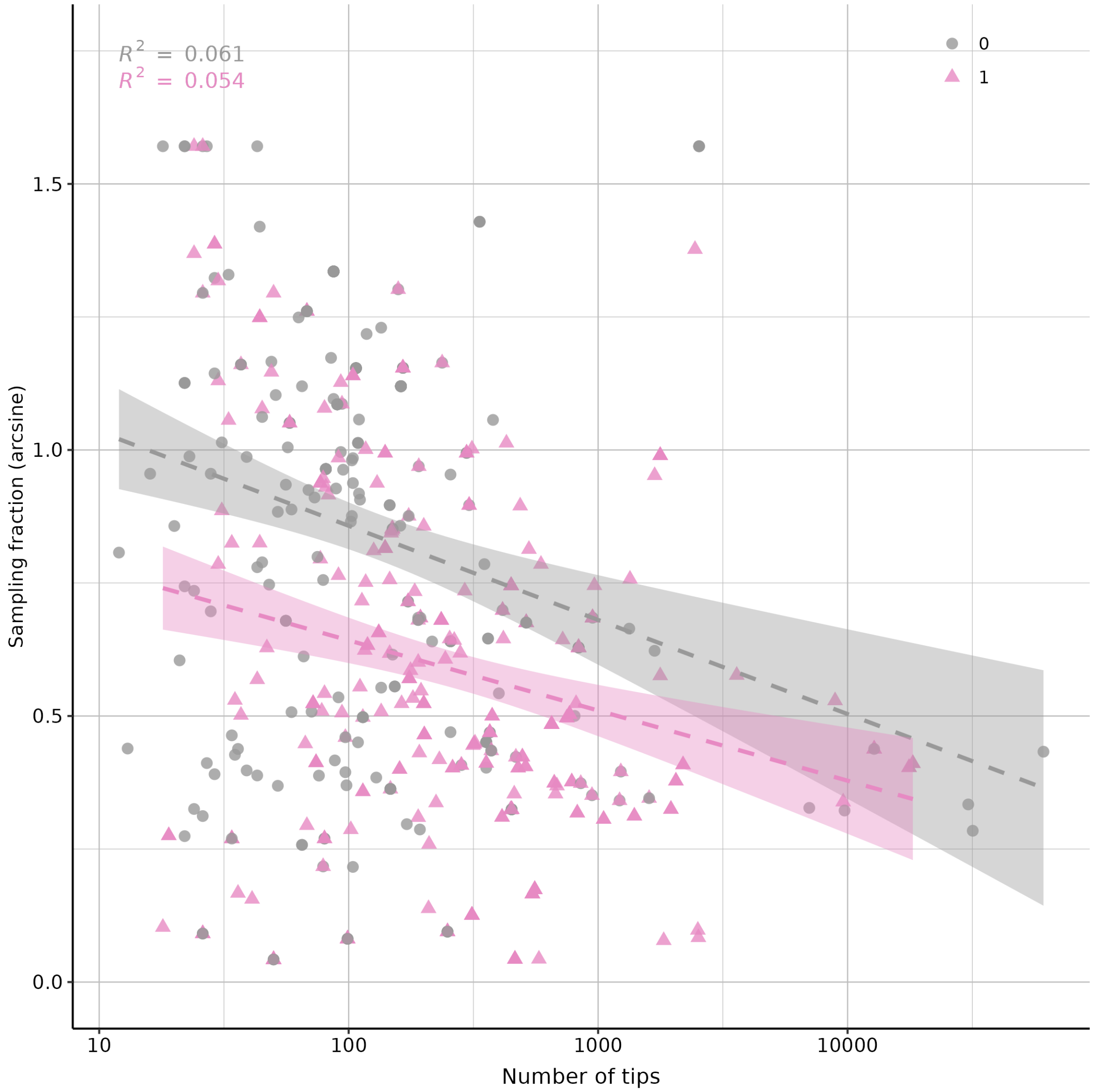


Figure S13: A scatterplot showing the relationship between sampling fraction of the tree used with an SSE model, and the number of tips in the tree. Coloured points are models for which trait-dependent diversification was detected, and grey points are models where it was not detected. Lines were fitted using linear models to these two groups with 95% confidence intervals estimated.

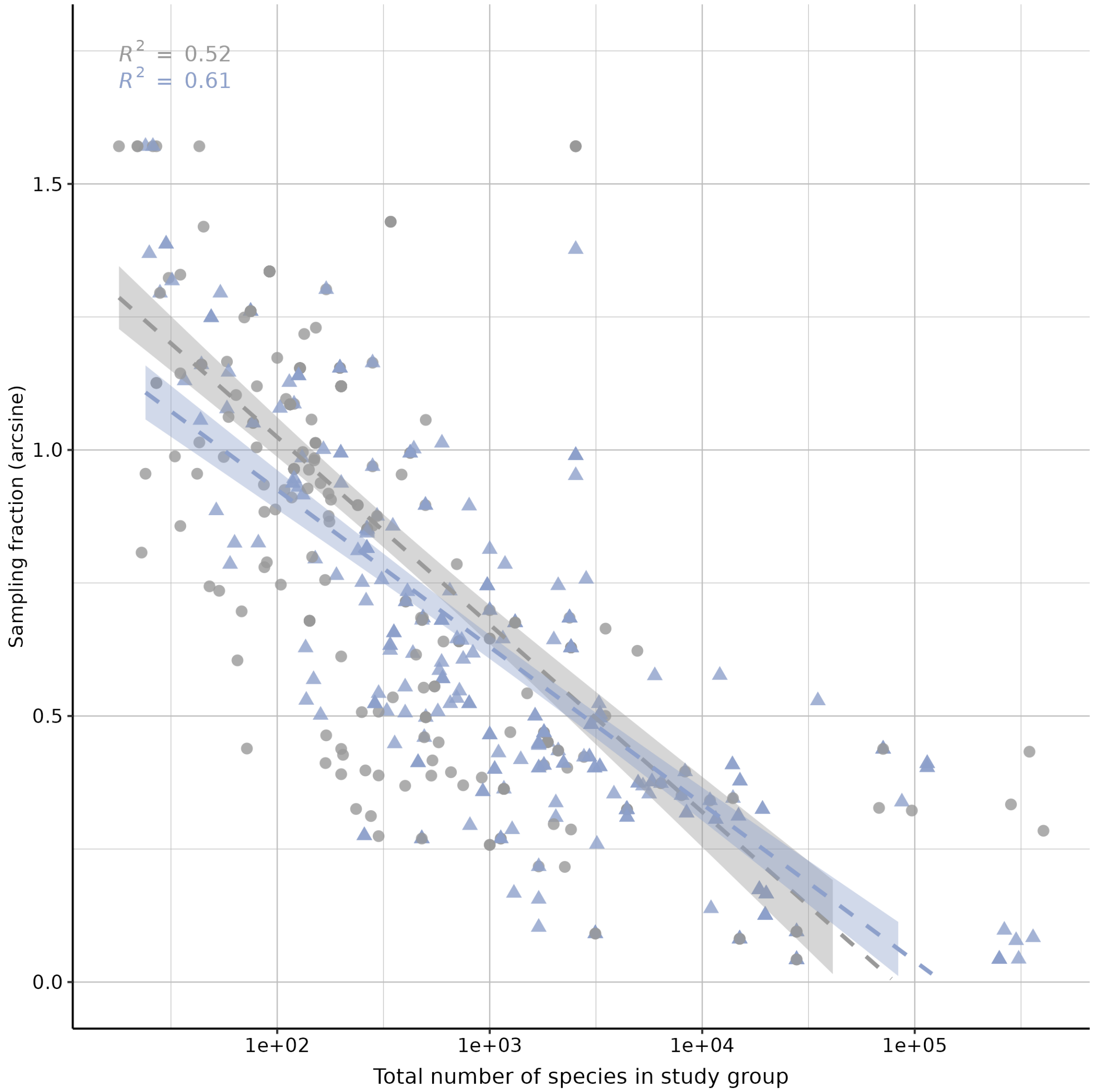


Figure S14: A scatterplot showing the relationship between sampling fraction of the tree used with an SSE model, and the total number of species in the study group the tree represents. Coloured points are models for which trait-dependent diversification was detected, and grey points are models where it was not detected. Lines were fitted using linear models to these two groups with 95% confidence intervals estimated.

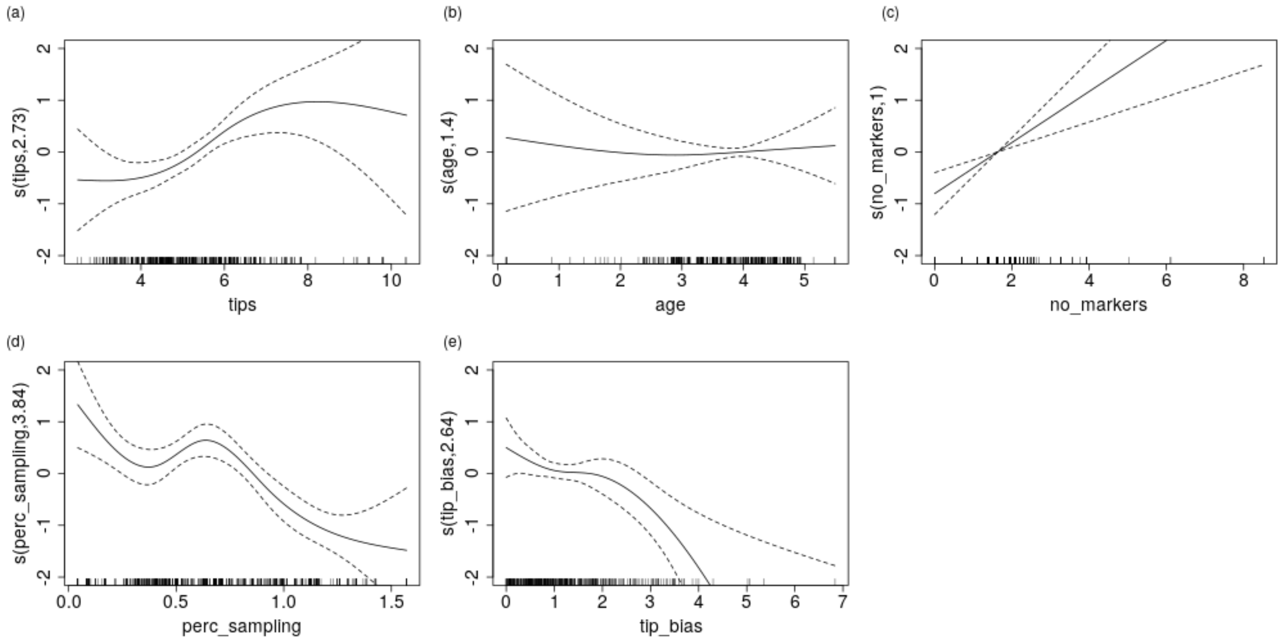


Figure S15: Panels (a-e) show the relationships in a generalized additive model that included all five of the continuous dataset properties in Figure 4 and SSE model outcome as the response variable. Missing values were replaced with column means. Values that are positive on the y-axis indicate that the dataset property at this x-axis value tends to be associated with trait-dependent diversification. Negative y-axis values indicate when dataset properties are associated with no effect. All variables were significant terms in the model except age, see Table S2 for full results.

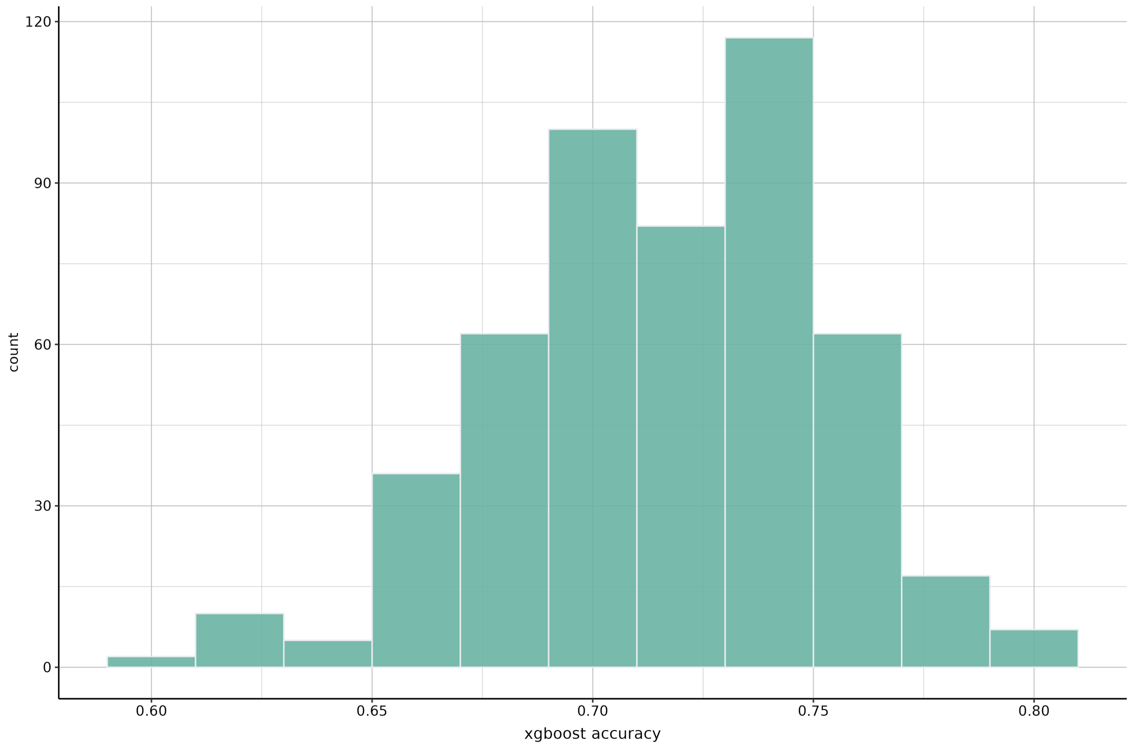


Figure S16: A histogram showing the distribution of accuracies (whether the correct SSE model outcome is predicted) after running 500 iterations of xgboosts. Each run used a different training (80%) and test (20%) datasets to capture the stochasticity in the dataset partitioning.

### 3 Supplementary tables

Table S1: A table showing the trait type ontology used to classify character states in state-dependent speciation and extinciton (SSE) models at six different levels. From left to right the classification becomes more specific. If a classification is not written at a given level, then the most specific classification that is written was used as the trait category (e.g. sexual system is used at level 5 and level 6).

| **Level 1** | **Level 2** | **Level 3** | **Level 4** | **Level 5** | **Level 6** |
|---|---|---|---|---|---|
| Extrinsic | Biogeography | Biome | | | |
| | | Region | | | |
| | Habitat | Soil | | | |
| | | Climate | | | |
| | | Elevation | | | |
| | | Vegetation | | | |
| | Elevation | | | | |
| Intrinsic | Vegetative | Growth | LifeSpan | | |
| | | | LifeForm | | |
| | | Morphology | PlantSize | | |
| | | | LeafMorpho | | |
| | | | PlantArchitecture | NrOfAxisCategories | |
| | | | MorphoOther | | |
| | | Physiology | Photosynthesis | | |
| | | | Fire | | |
| | | | Dormancy | | |
| | Reproduction | Pre-mating | BreedingSystem | SexualSystem | |
| | | | | MatingSystem | |
| | | | | SexAsex | |
| | | | FlowerMorpho | Inflorescence | |
| | | | | FlowerGeneral | FlowerSize |
| | | | | | FlowerSymmetry |
| | | | | | FlowerShape |
| | | | | | FlowerColor |
| | | | | | Reward |
| | | | | Male | Anthers |
| | | | | | AntherGlands |
| | | | | | Pollen |
| | | | | Female | Pistil |
| | | Post-mating | FruitMorpho | FruitSize | |
| | | | | FruitType | |
| | | | | FruitColor | |
| | | | SeedMorpho | SeedShape | |
| | | | | SeedSize | |
| | | | | SeedWings | |
| | Genome | Ploidy | | | |
| | | ChromosomeNumber | | | |
| Interactions | Defense | | | | |
| | Symbiosis | | | | |
| | Pollination | | | | |
| | Dispersal | | | | |

Table S2: Results of the full generalized additive model (GAM) including five continuous dataset properties with SSE model outcome (trait-dependent diversification vs no effect) as the response. 'edf' is the estimated degrees of freedom for the model terms. 'Ref.df' is the reference degrees of freedom. 'Chi.sq' is the test statistic for assessing the significance of model smooth term.

| **term** | **edf** | **Ref.df** | **Chi.sq** | **p-value** |
|---|---|---|---|---|
| s(tips) | 2.725 | 3.240 | 14.700 | 0.003 ** |
| s(age) | 1.396 | 1.685 | 0.480 | 0.785 |
| s(no_markers) | 1.000 | 1.000 | 15.750 | 7.22e-05 *** |
| s(perc_sampling) | 3.845 | 3.984 | 46.090 | <2e-16 *** |
| s(tip_bias) | 2.639 | 3.103 | 14.550 | 0.003 ** |
